# A pan-serotype human monoclonal antibody protects against pneumococcal infection by targeting multiple choline binding domain proteins

**DOI:** 10.64898/2026.05.28.728567

**Authors:** Anna L. McCormick, Behrouz Ghazi Esfahani, Caroline K. Page, Justin D. Shepard, Ana G. Jop Vidal, Katelyn D. McCaffrey, F. Eun-Hyung Lee, S. Mark Tompkins, Jorge E. Vidal, Jarrod J. Mousa

**Affiliations:** Department of Biomedical Sciences, College of Medicine, Florida State University, Tallahassee, FL, USA; Department of Infectious Diseases, College of Veterinary Medicine, University of Georgia, Athens, GA, USA; Department of Molecular Biosciences, Northwestern University, Evanston, IL, USA; Center for Vaccines and Immunology, College of Veterinary Medicine, University of Georgia, Athens, GA, USA; Department of Cell and Molecular Biology, and Center for Immunology and Microbial Research, School of Medicine, The University of Mississippi Medical Center, Jackson, MS, USA; Department of Medicine, Emory University, Atlanta, GA, USA

## Abstract

*Streptococcus pneumoniae* remains a global health threat, particularly to young children, the elderly, and immunocompromised individuals. Pneumococcal vaccines targeting the bacterial capsule polysaccharide do not protect against all 100+ pneumococcal serotypes, contributing to non-vaccine serotype infections and antibiotic resistance. To address these limitations, we isolated human monoclonal antibodies (mAbs) targeting pneumococcal surface proteins and identified a first-in-class mAb, derived from a patient with prior pneumococcal infection, namely mAb 5995-40. mAb 5995-40 bound multiple pneumococcal proteins, including PcpA and PspA, through a conserved choline-binding domain shared across serotypes. Functionally, mAb 5995-40 provided complete protection in lethal pneumococcal challenge models and improved survival in influenza A, influenza B, and respiratory syncytial virus-associated bacterial coinfection models. Mechanistic studies showed enhanced opsonophagocytic killing, reduced bacterial dissemination, and blocked epithelial translocation. Cryo-electron microscopy identified a repeating motif within the choline-binding domain targeted by mAb 5995-40, highlighting its potential as a broadly protective pneumococcal therapeutic.

## Introduction

*Streptococcus pneumoniae* (also known as the pneumococcus) is a gram-positive, extracellular opportunistic bacterial pathogen that remains a leading cause of morbidity and mortality despite the widespread use of multiple Food and Drug Administration (FDA)-approved vaccines.^1,2^ *S. pneumoniae* can cause mild infections, such as otitis and sinusitis, but can also disseminate into secondary organs to cause severe invasive pneumococcal disease, such as pneumonia, sepsis, and meningitis.^3^ The World Health Organization (WHO) estimates that over 1 million deaths occur worldwide each year due to pneumococcal infection.^4^ Children below the age of two, adults above the age of 65, and immunocompromised individuals are more susceptible to invasive pneumococcal disease.^1^

Currently, pneumococcal vaccination for disease prevention and antibiotics for disease treatment are widely used and have been highly successful at reducing the disease burden of *S. pneumoniae.*^5^ However, even with the use of protective vaccines and treatments, persistent rates of pneumococcal infection remain worldwide.^6^ Pneumococcal vaccines elicit antibodies against the capsular polysaccharide (CPS), which plays a critical role in virulence and evasion of the host immune response.^5,7^ However, over 100 different CPS serotypes exist and pneumococcal vaccines only cover a subset of them, with either a 23-valent pneumococcal polysaccharide vaccine (PPSV23) or multivalent pneumococcal conjugate vaccines (PCV15, PCV20, or PCV21) available. Due to low vaccination coverage of serotypes, there has been an increase in pneumococcal infections caused by non-vaccine serotypes and non-encapsulated strains, which remain unaffected by prior pneumococcal vaccination.^5,8,9^ Additionally, antibiotic resistance is common within non-vaccine serotypes, with some serotypes showing multi-drug resistance.^10^

These limitations have prompted investigation into alternative strategies, including the development of vaccines targeting broadly conserved surface proteins of *S. pneumoniae*. A significant area of research is therefore directed toward identifying and incorporating conserved surface proteins into pneumococcal vaccination that may provide broader and serotype-independent protection.^1,11^ Multiple pneumococcal antigens have been tested as vaccines in preclinical animal models, including the toxin pneumolysin, pneumococcal surface protein A (PspA), pneumococcal histidine triad protein D (PhtD), and pneumococcal surface adhesin A (PsaA), with several of these entering clinical trials.^12^ PspA is one of the most extensively studied choline-binding proteins and has been investigated for more than three decades as an immunogen and potential vaccine candidate.^13^ In 2024, a protein-based vaccine formulation containing recombinant PspA and the detoxified pneumolysin was evaluated in a Phase Ib clinical trial (NCT05622942).^14^ This study was shown to be safe and well-tolerated while demonstrating increased humoral immune responses at 30 days post-vaccination compared to PPSV23 in human participants.^15^ Similarly, PsaA has been assessed in a Phase I clinical trial as a part of a multicomponent protein-based vaccine formulated with two additional pneumococcal surface proteins: the serine/threonine-protein kinase P (StkP) and the protein required for cell separation (PcsB) (NCT00873431).^16^ This trial demonstrated both safety and immunogenicity against the included antigens.^17^

Another important immunogen, PhtD, is a member of the PHT family of conserved proteins on *S. pneumoniae* that all share histidine triad motifs.^18^ PhtD is immunogenic and induces protective anti-PhtD antibodies that reduce pneumococcal infection.^19^ Vaccination with PhtD has shown protection against lethal pneumococcal disease in a mouse model, and immunization with both PhtD and the detoxified pneumolysin led to increased protection against pneumococcal disease in rhesus macaques.^18,20,21^ PhtD, formulated with or without detoxified pneumolysin, was evaluated in a phase IIb clinical trial in which it was administered along with PCV13.^22^ The addition of PhtD did not demonstrate a significant improvement in protection compared to PCV13 alone. Because PhtD was not tested as a standalone vaccine, a direct assessment of its independent efficacy could not be performed. Nevertheless, the trial provided important insights into the potential role of conserved pneumococcal proteins in future protein-based vaccine formulations.^4^

Monoclonal antibodies (mAbs) are promising tools for broad treatment of pneumococcal infection independent of serotype while avoiding the potential complications of antibiotic resistance.^23^ We have previously isolated the first human mAbs targeting pneumococcal protein antigens, namely PhtD.^1^ Multiple mAbs targeting PhtD, specifically mAb PhtD3 and PhtD7, were shown to increase protection against lethal pneumococcal infection in a mouse model.^1^ The protective efficacy of mAb PhtD3 was dependent on macrophages and complement, as depletion of either component significantly reduced protection *in vivo.*^5^ *In vitro* analysis demonstrated that mAb PhtD3 enhanced opsonophagocytic uptake and killing of *S. pneumoniae* bacteria by differentiated HL-60 cells.^1^ Notably, a combination of mAbs PhtD3+PhtD7 provided increased protection in multiple mouse models of viral-bacterial co-infection, including respiratory syncytial virus (RSV), human metapneumovirus (hMPV), and influenza A virus.^24^

Since mAbs PhtD3 and PhtD7 are protective against both pneumococcal infection and viral co-infections, it is reasonable to assume that human mAbs targeting other conserved surface proteins would be protective. PcpA and PspA are both members of a group of surface proteins called choline binding proteins, which all share a non-conserved functional domain, conserved proline-rich region, and conserved choline-binding domain.^25^ Previous work has focused on the conserved proline-rich region, as it exhibits high sequence homology across serotypes.^26^ However, there is limited research on the conserved choline-binding domain, which consists of repeating structural motifs allowing for the non-covalent attachment to the phosphocholine moiety located within the bacterial cell wall.^27^ The choline-binding domain is a critical target because it mediated attachment of these choline binding proteins to the bacterial cell surface, where disruption of this interaction impairs bacterial infection and replication. In this study, we generated human mAbs against nine pneumococcal surface antigens from human subjects. While several mAbs had limited serotype breadth or lacked protection *in vivo*, one mAb, 5995-40, demonstrated extraordinarily high serotype breadth. mAb 5995-40 conferred protection in multiple animal models, including prophylactic and therapeutic treatments against two serotypes of *S. pneumoniae*. Functional studies revealed that the protective efficacy of mAb 5995-40 depends on complement activation, phagocytosis, and inhibition of bacterial translocation across the respiratory epithelium. Its efficacy was further evaluated in co-infection models, including influenza A, influenza B, and respiratory syncytial virus (RSV), followed by pneumococcal infection. Finally, cryo-electron microscopy (cryo-EM) showed that mAb 5995-40 cross-reacts with multiple choline binding proteins by targeting a conserved repeating unit within the choline-binding domain.

## Results

### Isolation of pneumococcal antigen-specific human mAbs

To identify pneumococcal-specific human mAbs, we recombinantly expressed his-tagged pneumococcal antigens that are well-conserved across serotypes, including PhtD, pneumococcal histidine triad protein E (PhtE)^28^, PspA, PsaA, PcpA, neuraminidase A (NanA)^29^, StkP, PcsB, and pneumococcal iron uptake protein A (PiuA)^30^, using sequences from either strain TCH8431 (serotype 19A) or strain TIGR4 (serotype 4). These proteins were used to isolate CD19^+^/IgD^-^/IgM^-^/IgA^-^ antigen-specific B cells **(Figure S1)** from human peripheral blood mononuclear cells through flow cytometry sorting. Sorted antigen-specific B cells were processed using the 10X chromium system followed by V(D)J library sequencing to obtain paired heavy and light chain variable region B cell receptor (BCR) sequences. From three separate experiments, 157 BCR sequences were obtained, and 110 corresponding mAbs were recombinantly expressed in HEK293F cells. The mAbs were screened against all recombinant pneumococcal proteins listed above by enzyme-linked immunosorbent assay (ELISA). Out of the 110 mAbs expressed, 21 mAbs bound to five different pneumococcal proteins, PhtE, PsaA, PcpA, NanA, PspA and PscB, at varying binding reactivities **(Figure 1A, Table S2, Figure S2)**. To determine the serotype breadth of the highest binding mAbs, we assessed mAb binding by flow cytometry (example flow panel in **Figure S3**) to eight different serotypes of *S. pneumoniae:* serotypes 1, 3, 4, 10A, 11A, 12A, 19A, and a multidrug- resistant serotype 3 (ABC02110133). Each mAb had variable binding across the panel of *S. pneumoniae* serotypes, with the lowest percentage of mAbs binding to serotype 3 **(Figure 1B)**.

**Figure 1:**
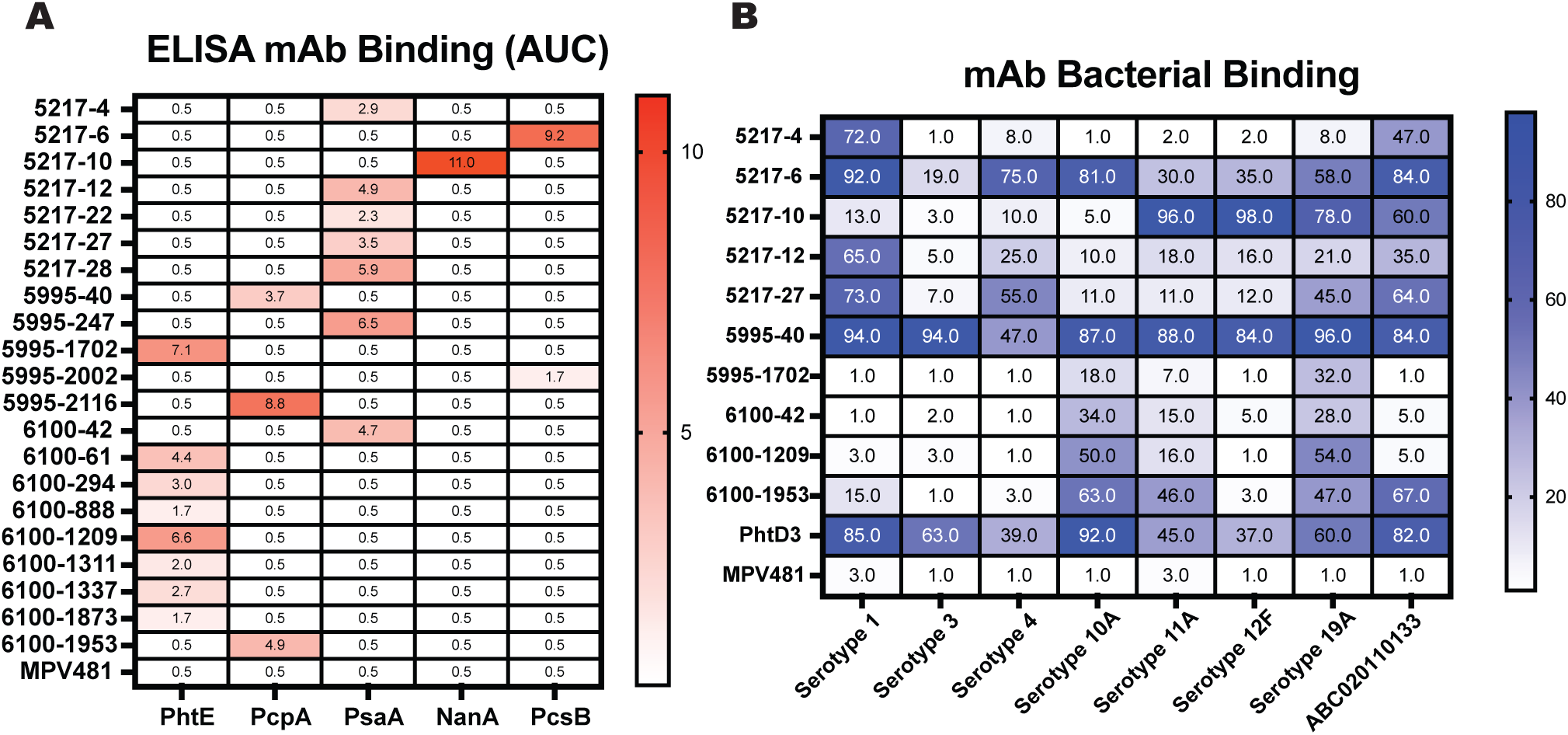
Binding profiles of purified mAbs to recombinant pneumococcal proteins and whole *S. pneumoniae* bacteria. (A) Binding of purified mAbs tested by ELISA against the 9 antigens included, with only 5 antigens showing mAb binding. Numbers in the box show the area under the curve for mAb binding, with the darker red color corresponding to higher binding. MPV481 was used as a negative control. (B) Binding of mAbs to a panel of clinically relevant whole fixed *S. pneumoniae* bacteria, with ABC02110133 being a multidrug resistant clinical isolate from the CDC. Numbers in the box show the percent binding of the mAb, with the darker blue color corresponding to higher binding. MPV481 was used as a negative control.

### Prophylactic treatment with pneumococcal mAbs leads to variable protection

Four pneumococcal mAbs (5217-6, 5217-10, 5995-1702, and 5995-40) targeting different antigens were selected for further *in vivo* analysis, as they demonstrated the broadest binding across pneumococcal serotypes, to determine whether prophylactic administration could protect mice against a lethal pneumococcal challenge. Mice were treated with the respective mAb 2 hours prior to infection at a dose of 15 mg/kg per mouse and then intranasally challenged with 1x10^7^ CFU/mouse of serotype 3 (WU2) *S. pneumoniae*. The infected mice were monitored over the course of 14 days for weight loss and the presence of clinical symptoms, such as labored breathing and lethargy. mAb 5217-6, which targets PcsB and demonstrated broad serotype binding **(Figure 1B)**, did not confer protection against pneumococcal challenge *in vivo* or reduce weight loss post infection **(Figure 2A, S4)**. Similarly, mAb 5217-10 (targeting PsaA) and mAb 5995-1702 (targeting PhtD), both of which exhibited more limited binding breadth **(Figure 1B)**, failed to provide protection or reduce weight loss following challenge **(Figure 2B, 2C, S4)**. In contrast, prophylactic treatment with mAb 5995-40 resulted in 100% survival compared to the PBS-treated control group, with minimal weight loss observed over the 14-day infection period **(Figure 3D, S4)**.

**Figure 2:**
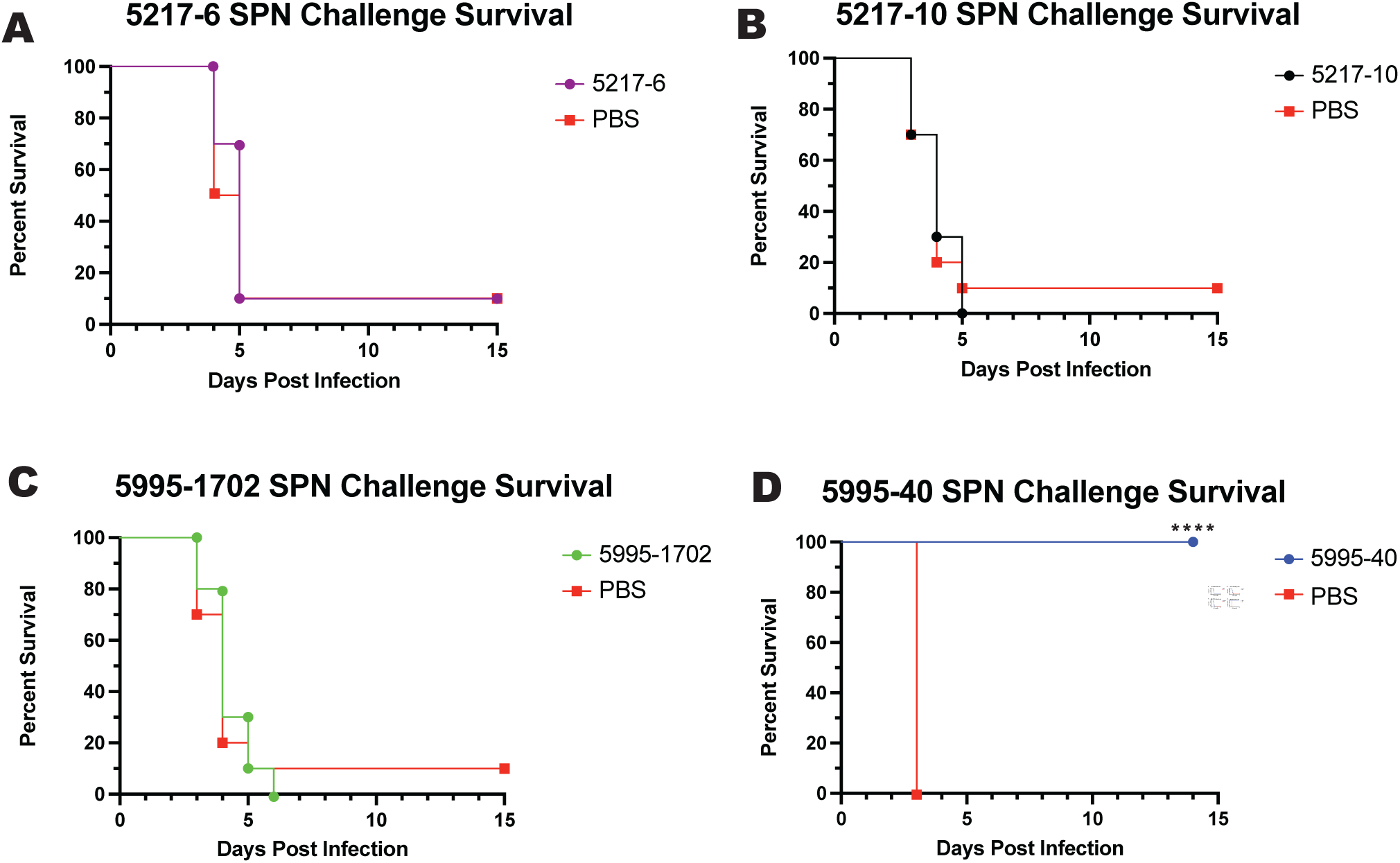
Determination of protection seen after lethal pneumococcal challenge. (A-D) Mice were intranasally challenged with 1x10^7^ CFU of WU2 (serotype 3) *S. pneumoniae* and treated prophylactically with the indicated mAb 2 hours prior to infection at a dose of 15 mg/kg per mouse. Body weight was monitored for 14 days post-infection, and survival was recorded. Percent survival is shown for each treatment group: (A) 5217-6 (purple), (B) 5217-10 (black), (C) 5995-1702 (green), and (D) 5995-40 (blue). **** p ≤ 0.001.

**Figure 3:**
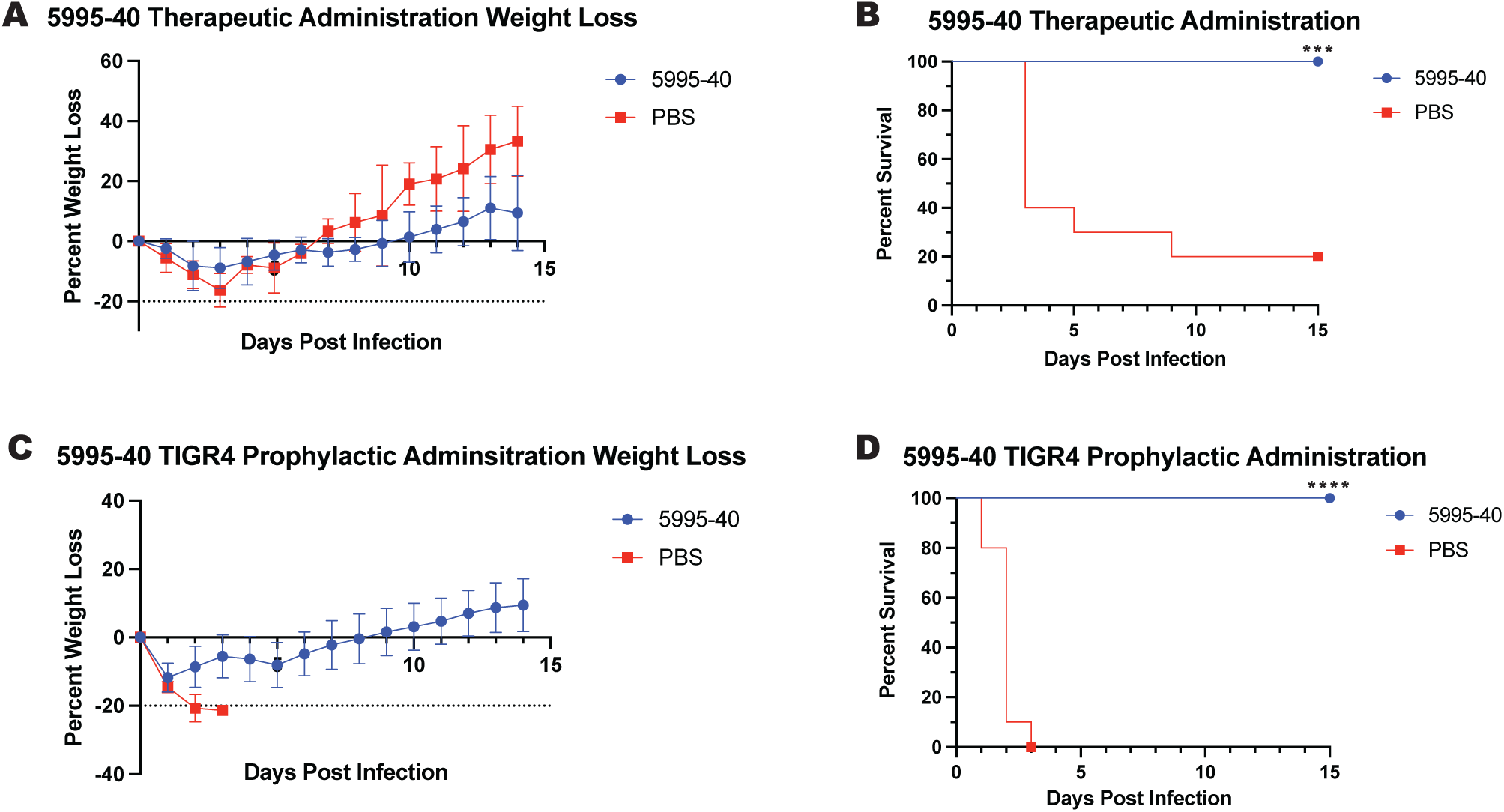
Further validation of mAb 5995-40 protection in lethal pneumococcal challenge models. (A-B) Mice were infected with 1x10^7^ CFU of WU2 (serotype 3) *S. pneumoniae* and treated 24 hours post-infection with mAb 5995-40 at a dose of 15 mg/kg per mouse. (A) Weight loss was measured over the course of 14 days, and (B) survival was recorded. (C-D) Mice were infected with 1x10^8^ CFU of TIGR4 (serotype 4) *S. pneumoniae* and treated prophylactically 2 hours prior to infection at a dose of 15 mg/kg per mouse. (C) Weight loss was measured over the course of 14 days, and (D) survival was recorded. Error bars are the standard deviation between living mice at each day post infection. ***, p ≤ 0.001; **** p ≤ 0.0001.

### Further validation of protective efficacy of mAb 5995-40 following both prophylactic and therapeutic administration

To further understand the protective capabilities of mAb 5995-40, an *in vivo* therapeutic treatment model was used. Mice were intranasally challenged with 1x10^7^ CFU/mouse of serotype 3 (WU2) *S. pneumoniae* and were treated with mAb 5995-40 24 hours post infection at a dose of 15 mg/kg per mouse. As seen in **Figure 3B**, treatment with mAb 5995-40 led to 100% survival when administered 24 hours post pneumococcal infection with minimal weight loss over the 14-day infection period (**Figure 3A)**, showing an increase in protection compared to the PBS-treated group. We also tested if mAb 5995-40 could protect against another pneumococcal serotype given its high binding breadth. To test this, an *in vivo* prophylactic treatment model using a lethal dose of TIGR4 (serotype 4) *S. pneumoniae* was used. Mice were intranasally challenged with 1x10^8^ CFU/mouse of TIGR4 and were treated with mAb 5995-40 2 hours prior to infection at a dose of 15 mg/kg per mouse. As shown in **Figure 3D**, treatment with mAb 5995-40 resulted in 100% survival compared to the PBS-treated controls, with minimal weight loss over the 14-day infection period (**Figure 3C)**. When considered alongside the serotype 3 (WU2) prophylactic and therapeutic studies, these findings demonstrate the broad protective capacity of mAb 5995-40, conferring robust protection across serotypes and independent of treatment timing. Based on these findings, mAb 5995-40 was selected for further investigation to elucidate the underlying mechanism of action.

### mAb 5995-40 binds to both PspA and PcpA within the conserved choline-binding domain

To better understand the breadth of binding of mAb 5995-40 across pneumococcal serotypes, an expanded panel of serotypes was used, including serotypes 1, 3, 4, 6A, 6B, 9V, 10A, 11A, 12F, 18C, 19A, 19B, 23C, and multidrug-resistant serotype 3 isolate from the Centers for Disease Control and Prevention (CDC). mAb 5995-40 showed binding across all serotypes tested **(Figure 4A)**. To further define the binding profile of mAb 5995-40, the mAb was tested against 9 pneumococcal proteins by ELISA , where binding was observed to 2 different proteins, PcpA and PspA, with similar binding patterns **(Figure 4B)**. To confirm these binding results, biolayer interferometry (BLI) was utilized to measure the binding rate of mAb 5995-40 to both PspA and PcpA. mAb 5995-40 bound to both full-length PspA and PcpA proteins, confirming the previous results **(Figure 4C)**. To better understand where mAb 5995-40 binds on these proteins, the full-length PspA protein was truncated into four different fragments: N-terminal domain, proline-rich region, C-terminal domain, and a combination of the N-terminal domain, and proline-rich region, and mAb 5995-40 was tested by ELISA to determine the binding to each fragment. mAb 5995-40 bound only to the C-terminal choline-binding domain (amino acids 436-725) **(Figure 4D)**. Both PspA and PcpA proteins contain a C-terminal choline-binding domain with 66% sequence homology between choline-binding domains, explaining the binding patterns seen between mAb 5995-40 and both antigens.

**Figure 4:**
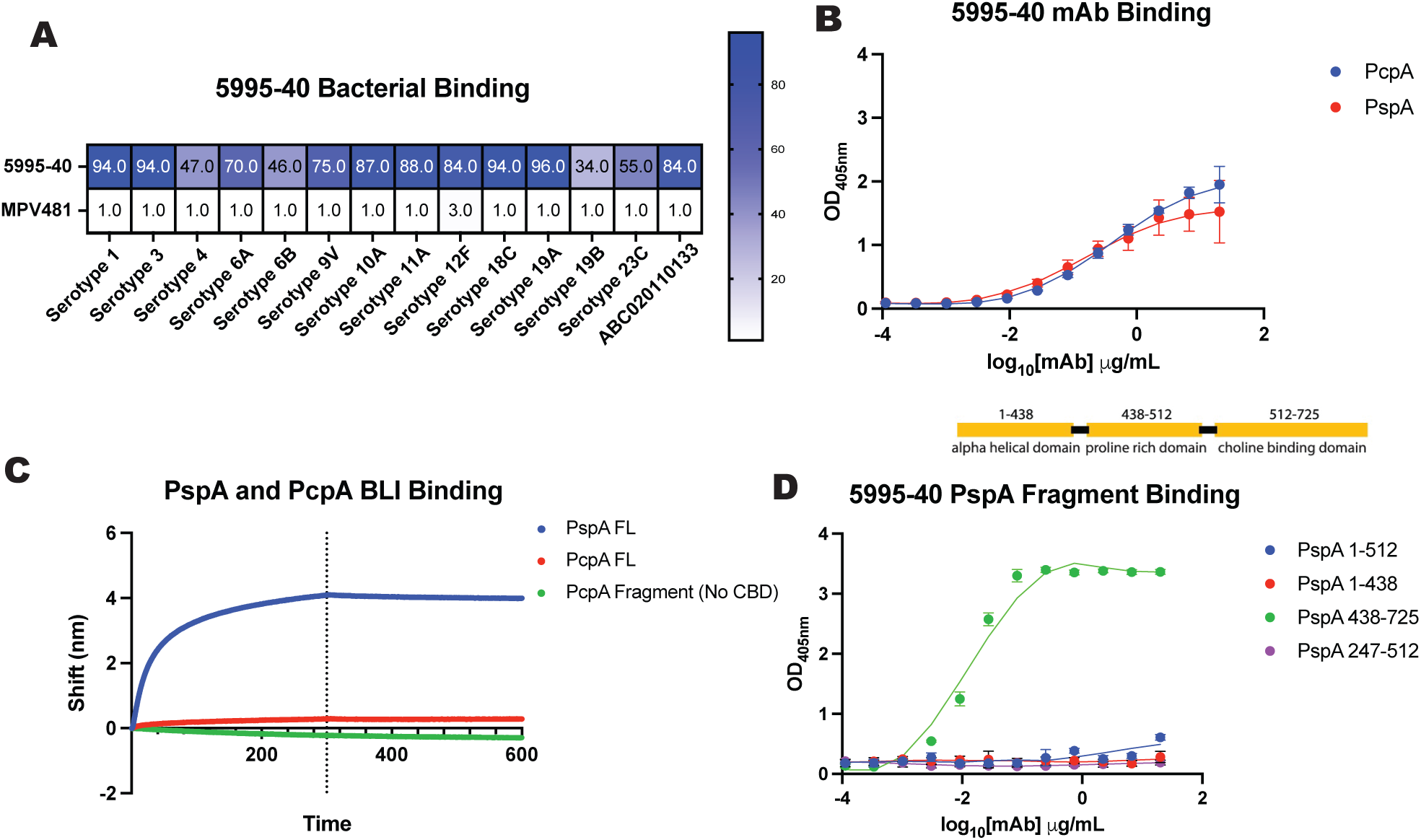
Complete binding profile of mAb 5995-40. (A) Binding of mAb 5995-40 mAb against expanded panel of *S. pneumoniae* serotypes, with ABC02110133 being a multidrug-resistant clinical isolate from the CDC. Numbers in the box show the percent binding of the mAb, with the darker blue color corresponding to higher binding. MPV481 was used as a negative control. (B) Binding of mAb 5995-40 to both PcpA and PspA recombinant proteins by ELISA, with each point on the graph correlating to the OD_450_ measured at that given mAb dilution. Error bars are the standard deviation between 4 technical replicates at each mAb dilution. (C) Binding of mAb 5995-40 to both PcpA FL and PspA FL recombinant proteins by BLI. PcpA fragment was used as a negative control, as the choline-binding domain was removed. (D) Binding of mAb 5995-40 to recombinant PspA fragments, with each point on the graph correlating to the OD_450_ measured at that given mAb dilution. Error bars are the standard deviation between 4 technical replicates at each mAb dilution.

### mAb 5995-40 binds to multiple proteins within the choline-binding protein family

*S. pneumoniae* has a family of 14 proteins called the choline-binding proteins, which all contain a C-terminal choline-binding domain that attaches the pneumococcal protein to the phosphocholine on the bacterial cell wall. These choline-binding domains are built of repeating amino acid motifs and share 30-99% sequence homology between the family of choline-binding proteins **(Figure 5A, 5B**). To determine if mAb 5995-40 could bind to multiple choline-binding proteins, mAb 5995-40 was tested by ELISA against 6 different choline-binding proteins: LytA, LytB, PcpA, PspA, PspC, and CbpE **(Figure 5C, S5)**. As shown in the heat map in **Figure 5C**, mAb 5995-40 showed high binding reactivity to all 6 choline-binding proteins, with the darker blue color corresponding to higher binding. To confirm these findings, BLI was utilized to test binding to multiple choline-binding proteins, including both full length proteins and the truncated choline binding domain. mAb 5995-40 bound with strong affinity to all choline-binding proteins tested as can be seen by the strong association of mAb 5995-40 to each protein with very little dissociation, as seen in **Figure 5D**.

**Figure 5:**
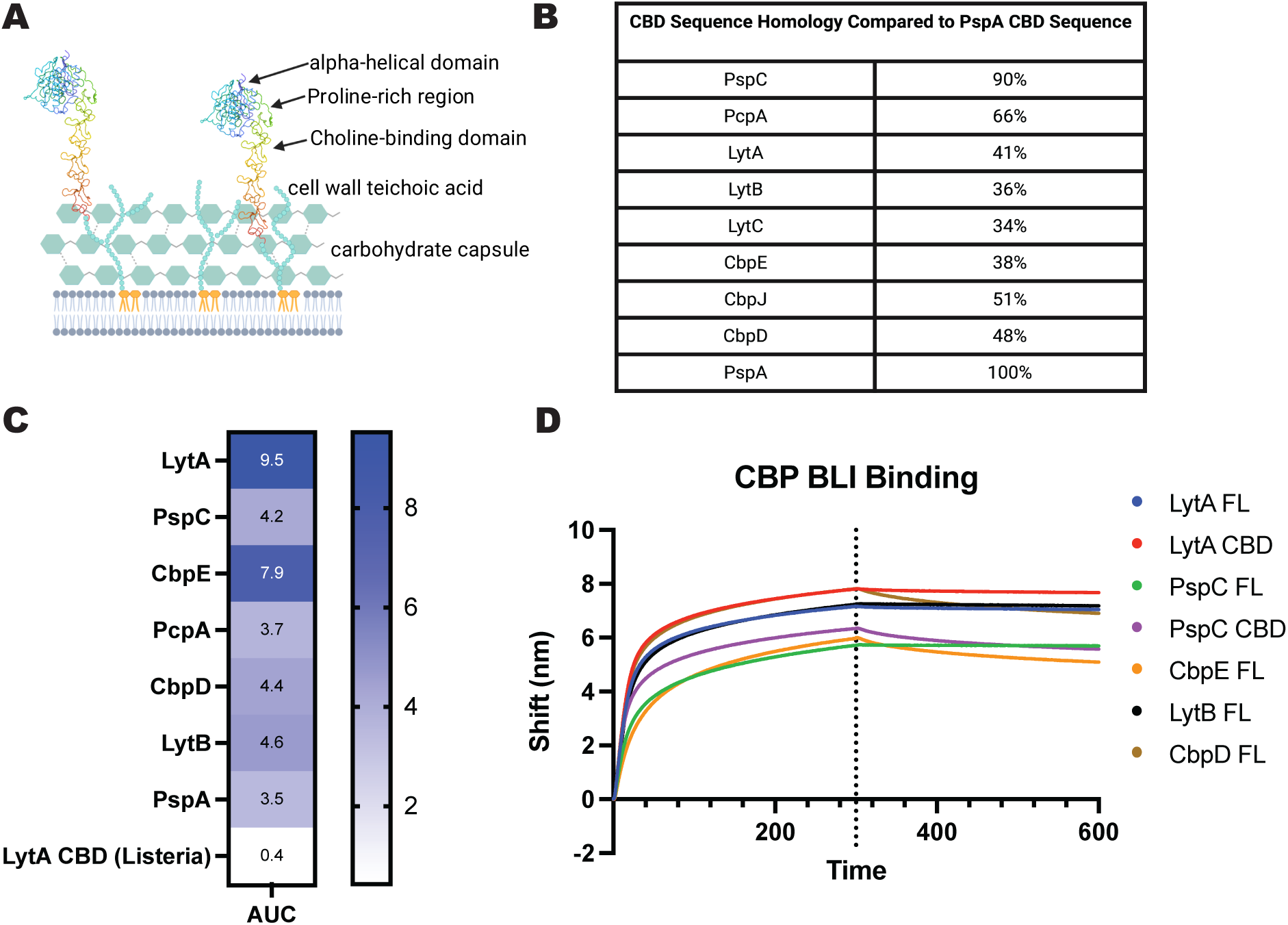
Binding breadth of mAb 5995-40 against family of choline-binding proteins on the surface of *S. pneumoniae*. (A) Schematic diagram of choline-binding protein on the surface of *S. pneumoniae*, containing the N-terminal functional domain, proline-rich region, and the C-terminal choline-binding domain. (B) The choline-binding domain (CBD) of each protein was aligned against the PspA CBD to determine percent sequence homology. Because the number of repeats varies among proteins, alignments were performed relative to the shorter CBD. (C) Heat map of mAb 5995-40 binding multiple choline-binding proteins by ELISA. Numbers in the box show the area under the curve for mAb binding, with the darker blue color corresponding to higher binding. (D) BLI binding for mAb 5995-40 binding to various CBPs, both full length and CBD.

### Treatment with mAb 5995-40 increases complement activation and pneumococcal bacterial uptake

Treatment with mAbs can recruit complement proteins in circulation to bind to the Fc region through the classical pathway.^31^ This process can result in the formation of the membrane attack complex or enhance opsonophagocytic clearance by innate immune cells.^31,32^ To assess the ability of mAb 5995-40 to promote complement activation, complement deposition was measured following mAb 5995-40 binding to two choline-binding proteins, PspC and LytA proteins, by flow cytometry. The addition of mAb 5995-40 increases the amount of C3 protein deposited on the surface of the antigen-coated beads, suggesting an increase in complement activation following treatment with mAb 5995-40 **(Figure 6A, 6B, S6)**. In addition to complement activation, treatment with mAbs can promote opsonophagocytic clearance through Fc-mediated engagement of phagocytic receptors.^33^ Antibody coating of the bacteria enhances their recognition and uptake by innate immune cells, including macrophages and neutrophils, thereby facilitating bacterial clearance.^33,34^ To evaluate whether mAb 5995-40 enhances this process, opsonophagocytic assays were performed to quantify the pneumococcal uptake by phagocytic cells **(Figure 6C, 6D).** The addition of mAb 5995-40 led to an increase in bacterial killing by differentiated HL-60 cells in comparison to no mAb, however, the levels of bacterial uptake were lower than the addition of serotype 4 specific serum, or with 007sp (PCV13 reference serum). This finding was the same for both wildtype TIGR4 **(Figure 6C)** and non-encapsulated TIGR4 (no CPS) **(Figure 6D)**. These data begin to provide a clearer picture of the mechanism of action of mAb 5995-40.

**Figure 6:**
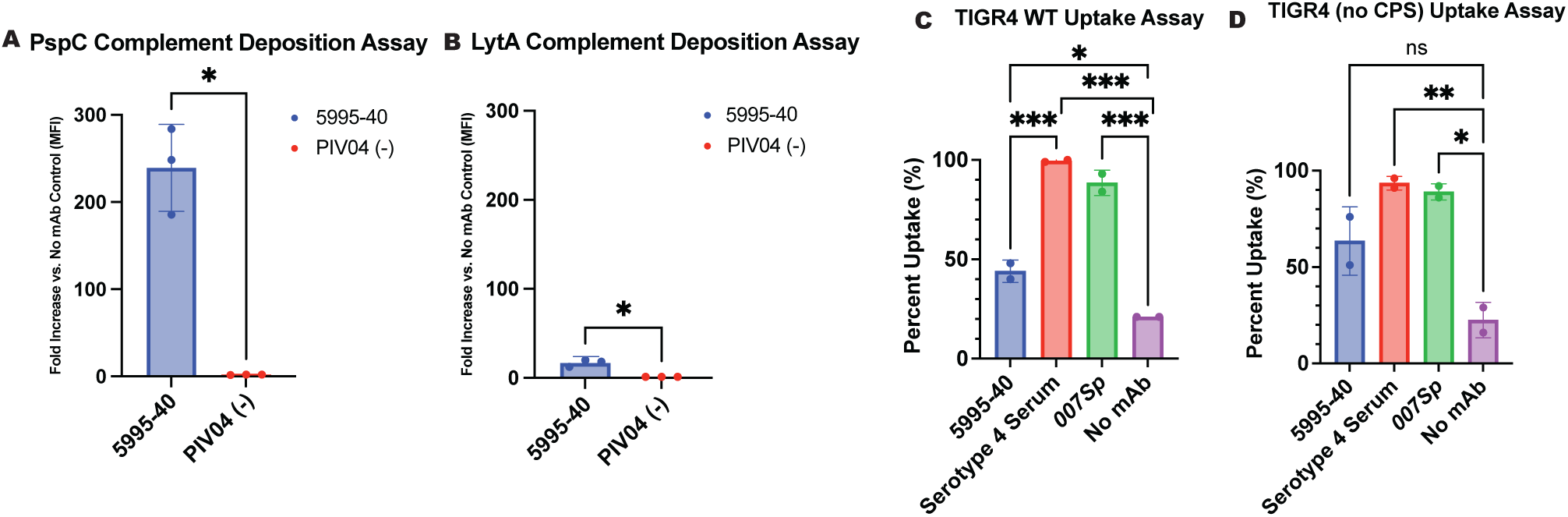
Determination of mechanistic properties of mAb 5995-40 through *in vitro* assays. (A-B) Antigen-coated beads were stained, and deposition of the C3b protein to the antigen-coated beads was measured. Complement deposition was tested against the proteins (A) PspC and (B) LytA, where fold increase in comparison to the no mAb control was graphed. Error bars are the standard deviation between three technical replicates. (C-D) Uptake assays were completed against (C) TIGR4 WT and (D) TIGR4 (no CPS), determining the percentage of uptake and killing of bacteria in the presence of mAb 5995-40, serotype 4 serum, 007Sp (PCV13 reference serum), and no mAb control. ns, non-significant; *, p ≤ 0.05, ** p ≤ 0.01; *** p ≤ 0.001.

### Treatment with mAb 5995-40 protects the epithelial barrier against pneumococcal translocation leading to a decrease in bacterial dissemination

To further elucidate the mechanism of action of mAb 5995-40, pneumococcal translocation across an epithelial barrier was evaluated. *S. pneumoniae* is capable of transversing the respiratory epithelium, facilitating dissemination within the host.^35^ To determine whether mAb 5995-40 could limit this process, an *in vitro* transwell system was established in which human bronchial epithelial cell monolayers were cultured on permeable supports, and bacteria were pre-treated with mAb 5995-40 before introduction to the epithelial cell monolayer to assess the epithelial translocation. The addition of mAb 5995-40 led to a significant decrease in bacterial translocation 4 hours post-infection, while there was no significant difference at any other time points **(Figure 7A)**. These data suggest that mAb 5995-40 can block the bacteria from being able to cross the lung epithelial layer during infection. To further test this hypothesis, the bacterial burden after treatment with mAb 5995-40 was examined. Mice were treated prophylactically with mAb 5995-40 2 hours prior to WU2 pneumococcal infection, and lung and blood bacterial titers were collected 24, 48, and 72 hours post-infection. As seen in **Figure 7B and 7C**, treatment with mAb 5995-40 decreased lung **(Figure 7B)** and blood **(Figure 7C)** bacterial titers at 48 and 72 hours post-infection, with no significant difference seen at 24 hours in either the lung or blood. Collectively, these data demonstrate that prophylactic administration of mAb 5995-40 enhances protection against pneumococcal challenge. This protection could be mediated by the inhibition of the localization of these choline-binding proteins to the surface of the bacteria, decreasing the virulence and replication and in turn, decreasing the spread of bacteria into the lungs. This protection may also be mediated, at least in part, by the inhibition of bacterial translocation across the respiratory epithelial barrier, where bacteria in the lung alveoli are unable to cross the epithelium and disseminate into the blood following infection.

**Figure 7:**
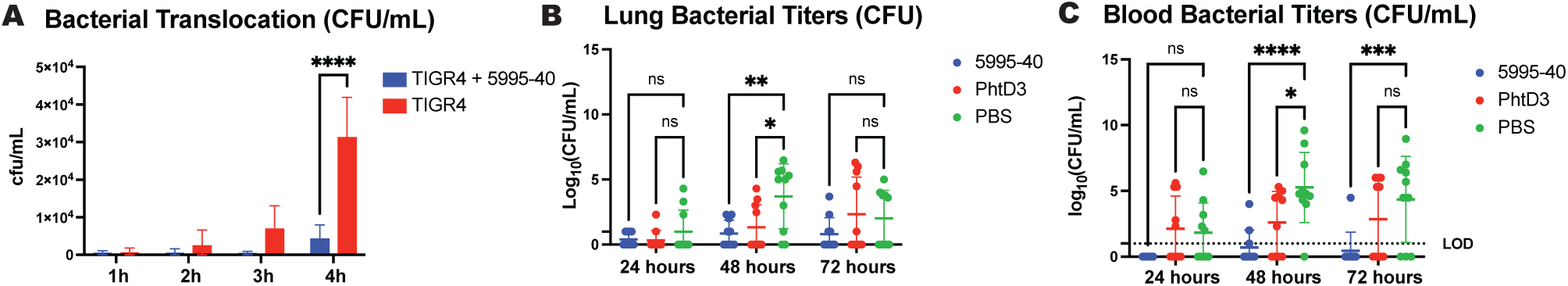
Assessment of mAb 5995-40 mediated inhibition of pneumococcal translocation. (A) Transwell plates coated with human bronchial epithelial cells were subject to infection with pre-treated pneumococcal bacteria, where the number of bacteria per mL (cfu/mL) is graphed for each time point tested. (B-C) Mice were challenged with WU2 (serotype 3) *S. pneumoniae,* and (B) lung and (C) blood bacterial titers were performed at 24, 48, and 72 hours post-infection. (n=10 mice per group). ns, non-significant; *, p ≤ 0.05, ** p ≤ 0.01; ***, p ≤ 0.001; **** p ≤ 0.0001.

### Prophylactic treatment with mAb 5995-40 protects against lethal viral-pneumococcal co-infections

Prior viral infection can dysregulate the host immune system, impairing the host’s antibacterial function and enhancing secondary bacterial infection^24^. Due to these virus mediated effects, we next determined if treatment with mAb 5995-40 could demonstrate similar protection upon secondary bacterial infection. Three viral-pneumococcal co-infection models were used: Influenza A (IAV/SPN), Influenza B (IBV/SPN), and RSV (RSV/SPN) all co-infected with serotype 3 (WU2) of *S. pneumoniae*. To access protection against the IAV/SPN co-infection, mice were infected with 100 PFU/mouse of A/California/07/2009 and infected again with 1x10^6^ CFU/mouse 7 days post-influenza infection. Mice were treated with 15 mg/kg of mAb 5995-40 2 hours prior to pneumococcal infection and were monitored for 14 days for weight loss and the presence of labored breathing, lethargy, and conjunctivitis of the eye. As seen in **Figure 8A**, mAb 5995-40 significantly increased protection against lethal IAV/SPN co-infection compared to the PBS negative control. To further investigate the protective effects of treatment with mAb 5995-40 against the IAV/SPN co-infection, lung and blood bacterial titers were collected 48 and 72 hours post-pneumococcal infection. Mice treated with mAb 5995-40 had a reduction in both lung and blood bacterial titers at all time points **(Figure 8B and 8C)**. Similarly, to assess protection against the IBV/SPN co-infection, mice were infected with 100 PFU/mouse B/Washington/02/2019, infected again with 1x10^6^ CFU/mouse 7 days post influenza infection, and treated and monitored as above. mAb 5995-40 significantly increased protection against a lethal IBV/SPN co-infection compared to the PBS negative control **(Figure 8D)**. From a separate group of mice, lung and blood bacterial titers were collected 48 and 72 hours post-pneumococcal infection, and mice treated with mAb 5995-40 had a significant reduction in both lung and blood bacterial titers at all time points **(Figure 8E and 8F)**. Lastly, to access protection against RSV/SPN co-infection, mice were infected with 1.7x10^6^ PFU/mouse of RSV A2 and subsequently infected with 1x10^6^ CFU/mouse 5 days post RSV infection. As above, mice were treated with 15 mg/kg of mAb 5995-40 2 hours prior to pneumococcal infection and were monitored for 14 days for weight loss and the presence of labored breathing and lethargy. As seen in **Figure 8G**, mAb 5995-40 significantly increased protection against a lethal RSV/SPN co-infection compared to the PBS negative control. We did not observe a significant reduction in lung or blood bacterial titers at 48 or 72 hours post- infection in mice treated with mAb 5995-40 **(Figure 8H and 8I)**.

**Figure 8:**
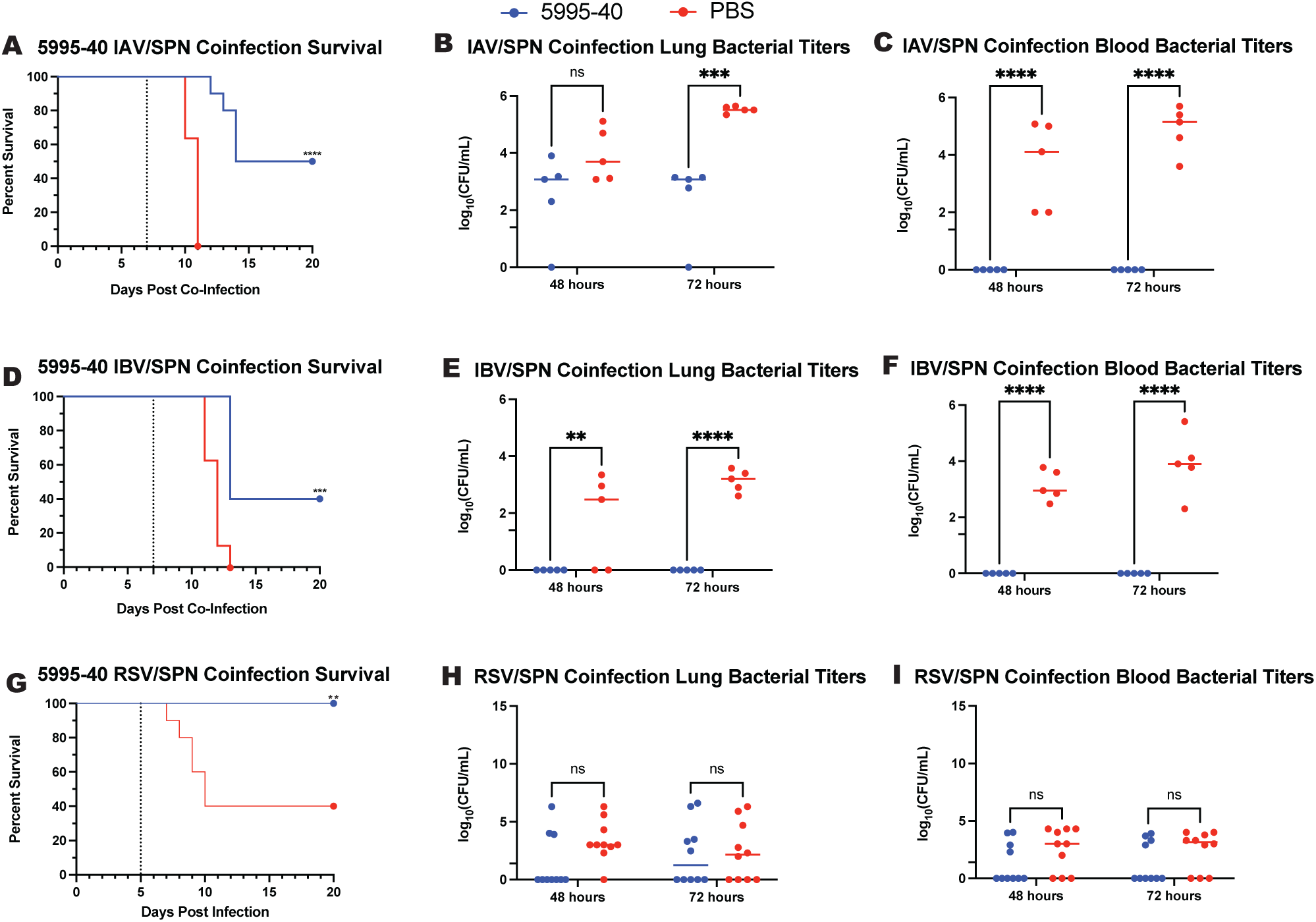
Determination of protection seen after lethal viral and pneumococcal co-infections. (A-C) Mice were challenged with a lethal co-infection of Influenza A and *S. pneumoniae.* Mice were administered mAb 5995-40 mAb 2 hours prior to *S. pneumoniae* infection. (A) Percent survival was calculated for each group over the course of 14 days, with 20% weight loss being the humane euthanasia point (n=10 mice per group). (B) Lung and (C) blood bacterial titers were performed at 48 and 72 hours post-pneumococcal infection (n=5 mice per group). (D-F) Mice were challenged with a lethal co-infection of Influenza B and *S. pneumoniae.* Mice were administered mAb 5995-40 mAb 2 hours prior to *S. pneumoniae* infection. (D) Percent survival was calculated for each group over the course of 14 days, with 20% weight loss being the humane euthanasia point (n=10 mice per group). (E) Lung and (F) blood bacterial titers were performed at 48 and 72 hours post-pneumococcal infection (n=5 mice per group). (G-I) Mice were challenged with a lethal coinfection of RSV and *S. pneumoniae*. Mice were administered mAb5995-40 mAb 2 hours prior to *S. pneumoniae* infection. (G) Percent survival was calculated for each group over the course of 14 days, with 30% weight loss being the humane euthanasia point (n=10 mice per group). (H) Lung and (I) blood bacterial titers were performed at 48 and 72 hours post-infection (n=10 mice per group). ns, non-significant; **, p ≤ 0.01; ***, p ≤ 0.001; ****, p ≤ 0.001

### mAb 5995-40 complexed with CbpE, a protein in the choline-binding protein family, demonstrates targeting of a conserved binding motif repeat

mAb 5995-40 was cleaved to Fab regions which were complexed with the full length CbpE protein, due to the high binding of mAb 5995-40 to CbpE and its large size. To determine the binding site of mAb 5995-40, we utilized cryo-electron microscopy (cryo-EM). After particle picking and heterogeneous refinement, we obtained an electron density map containing the full length CbpE protein with the 5995-40 Fab bound that was further processed to a global resolution of 3.4 Å **(Table S1, Figure S7)**. We leveraged Alphafold3 to predict a structure of the CbpE and 5995-40 structures separately and then used the CryoEM map to better fit the models and interactions. Models reveal the role of the CDRH1 and CDRH3 of the 5995-40 Fab in binding to the CbpE. mAb 5995-40 primarily uses HCDR1 and HCDR3 to interact with the CbpE protein **(Figure 9A)**. Based on our experimentally determined structure, there are two hydrogen bonds between the 5995-40 Fab and the choline-binding domain on the CbpE protein, one occurring between the LC CDR1 (W37 with K314) of 5995-40 Fab and the other through the HC CDR3 (T112 with W312). Another interaction is the HC CDR3 (P114) into a negatively charged electrostatic pocket consisting of SER369 and ASP368. The choline-binding domain of the pneumococcal choline-binding proteins is composed of repeating amino acid motifs that form a coiled structure responsible for anchoring these proteins to the bacterial cell wall. Our structural analysis also revealed that the 5995-40 Fab binds multiple sites along the choline-binding domain **(Figure 9C, S8)**. Sequence examination further identified repeats of tryptophan (W) to lysine (K) residues 2 amino acids apart from each other, forming the repeating *WxK* motif, as shown in the black boxes in **Figure 9B**. These repeating motifs likely provide recurring structural elements that facilitate the hydrogen bonds shown in **Figure 9A** and stabilize Fab binding across multiple epitopes within the domain. Consistent with this model, up to three Fab molecules were observed to bind to a single CbpE protein simultaneously **(Figure 9C, S8).** Together, these findings indicate that the repetitive nature of the choline-binding domain enables multivalent engagement by mAb 5995-40, providing a structural basis for its broad reactivity across these choline-binding proteins.

**Figure 9:**
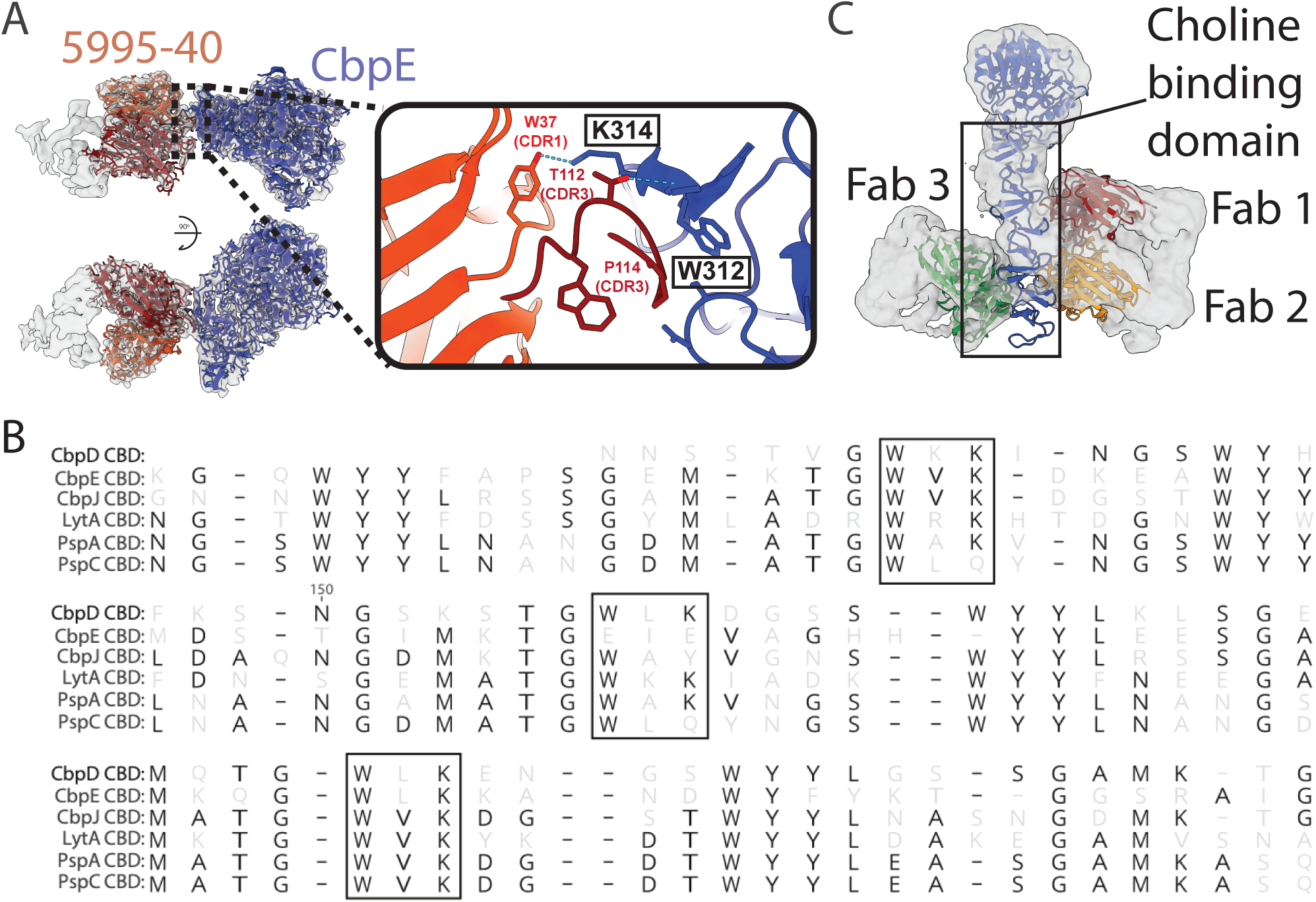
Cryo-EM map of 5995-40 Fab complexed with CbpE protein, a member of the choline-binding domain. (A) Model fit in the CryoEM map showing the overall and molecular level binding of the 5992-2 Fab (heavy chain: red, light chain: orange) to the CbpE antigen (navy). Interactions of the 5995-40 CDR1 and CDR3 with the CbpE protein in the choline-binding domain are marked by two hydrogen bonds indicated with light blue dotted lines. 5995-40 Fab (red) and CbpE protein residues associated with binding are labeled. (B) Sequence alignment of partial CBD sequences demonstrating some of the repeating *WxK* motifs in black boxes. (C) Overlay of the Alphafold3 predicted structure and experimentally determined structure, showing multiple 5995-40 Fabs binding to the C-terminal choline-binding domain of the CbpE protein.

## Discussion

In this study, we isolated and determined the binding affinity, serotype breadth, and protective properties of the first human mAb to target the choline-binding domain across pneumococcal choline-binding proteins. The choline-binding protein family contains many immunogenic surface proteins that have been examined in depth as vaccine candidates, including both PspA and PcpA, although the current outlook for progress of these antigens in the era of conjugated vaccines remains uncertain.^36,37^ However, human mAbs targeting these conserved antigens offer broader recognition across pneumococcal serotypes and potential for disease prevention and treatment independent of serotype.

Although there is great potential for human mAbs against *S. pneumoniae,* it is still largely understudied. This study aimed to provide a better understanding of immunogenic surface proteins after colonization or infection with *S. pneumoniae,* as 9 different SPN surface proteins were tested and only 5 showed binding to isolated mAbs. However, even if these mAbs bound to their respective proteins by ELISA, the true correlation to protection was dependent on the bacterial binding. Even though the antigens included in the study are known to be conserved across serotypes, there are varying binding patterns for each mAb across serotype, with most mAbs binding strongly to the serotype the antigen was cloned from and weaker to serotype 3. This would be expected as serotype 3 has a “sticky” capsule that is easily shed during infection, providing an added challenge while trying to target and treat.^38^ This pattern was evident for mAbs 5217-6, 5217-10, and 5995-1702, which, despite strong ELISA reactivity to their respective antigen, displayed low to moderate binding to serotype 3 bacteria and further failed to confer protection in a lethal serotype 3 challenge model. In contrast, mAb 5995-40 maintained robust binding across all tested serotypes, including serotype 3, and further provided significant protection in both serotype 3 and serotype 4 lethal infection models. Serotype 3 is a well-known vaccine-escape serotype^39^, while serotype 4 infections have recently reemerged in developed countries, highlight the potential of mAb 5995-40 as a broadly protective therapeutic candidate capable of addressing key gaps in current pneumococcal vaccination strategies against persistent and re-emerging serotypes^40,41^.

While mAb 5995-40 bound strongly across all serotypes of SPN, our subsequent work showed the mAb binding to two pneumococcal antigens, PcpA and PspA. Monoclonal antibodies are known to be extremely specific and typically bind to a single distinct epitope^42^, however, mAb 5995-40 seemed to bind to multiple. To better understand where mAb 5995-40 bound, a fragmented PspA protein was used, containing the N-terminal functional domain, the conserved proline-rich region, and the C-terminal choline-binding domain, where it was clear that mAb 5995-40 bound to the choline-binding domain, which is found in both PspA and PcpA. While the choline-binding protein family on *S. pneumoniae* is well studied, most work has focused on the non-conserved N-terminal functional domain or conserved proline-rich region for development of therapeutics.^36^ The choline-binding domain as a therapeutic remains largely understudied, as the choline-binding protein family share between 30-99% sequence homology. While looking into the sequences of each choline-binding domain, there were multiple repeat motifs found within each choline-binding protein sequence that could explain the binding patterns seen, where mAb 5995-40 is able to bind to multiple choline-binding proteins with strong affinities.

Anti-pneumococcal mAbs have potential for use in the clinic, as current vaccines cover only a subset of current serotypes.^1^ The prophylactic efficacy of mAb 5995-40 was established against pneumococcal serotype 3, a leading cause of invasive pneumococcal disease. We have demonstrated that mAb 5995-40 significantly improved survival when administered both prophylactically (2 hours prior to infection) and therapeutically (24 hours post-infection). In addition, prophylactic treatment with mAb 5995-40 enhanced survival following challenge with the TIGR4, serotype 4, strain of *S. pneumoniae*. Collectively, these findings indicate that mAb 5995-40 provides broad protection across multiple pneumococcal serotypes, independent of treatment timing. Consistent with the improved survival, treatment with mAb 5995-40 was also associated with reduction of bacterial burden in the lungs and blood at 24, 48, and 72 hours post-infection. These data show the reduced blood bacterial titers align with the decrease in translocation of the pneumococcus across the epithelial barrier *in vitro*. These data together suggest that mAb 5995-40 improves the host’s barrier function during pneumococcal infection to reduce the systemic spread of infection throughout the body.

Bacterial co-infections with viruses represent a significant challenge to public health.^43^ Primary influenza virus infection has been shown to predispose hosts to synergistic secondary bacterial infections leading to increased morbidity and mortality.^43^ From previous literature, it is known that influenza lung damage is usually greatest on day 6 post-infection,^44^ with the greatest susceptibility to secondary bacterial infection occurring near day 7 in humans and mice.^45^ With this information, we utilized a co-infection model that started the secondary pneumococcal infection 7 days post-influenza infection, with both an influenza A and influenza B primary viral infection. In both models, the survival protection of mAb 5995-40 was reduced compared to the protective efficacy observed in primary infection studies, however, there were little to no detectable bacterial titers in the lung and blood 48 and 72 hours post infection. Further studies are needed to decipher the exact protective mechanism in the context of IAV/SPN and IBV/SPN co-infections.

RSV, a member of the *Pneumoviridae* family, is a second major viral cause of secondary bacterial co-infections. Previous work found that in RSV/SPN co-infections, day 5 post viral provided the greatest susceptibility for secondary pneumococcal infection.^24^ In this RSV/SPN co-infection model, mAb 5995-40 provided 100% protection, similar to the protective efficacy during primary infections. However, it was seen that there was no difference between lung or blood bacterial titers at 48 or 72 hours post-pneumococcal infection. These data may be explained by the interactions between RSV and SPN during co-infection where it is known that RSV and SPN will create RSV-SPN complexes, expressing the viral glycoproteins on the surface of infected cells.^46,47^ This can lead to increased attachment and adherence, causing the dissemination of bacteria to occur more readily independent of treatment.^46,47^

In addition to the mechanistic and efficacy studies, we resolved the structure of mAb 5995-40 bound to the choline-binding domain of the pneumococcal protein, CbpE. Structural analysis revealed that mAb 5995-40 stabilizes its interaction through hydrogen bonding to two conserved amino acids within the repeating motifs of the choline-binding domain, enabling multivalent engagement along the length of this domain. Although monoclonal antibodies typically recognize a single discrete epitope, this epitope is reiterated multiple times within the choline-binding domain, allowing several 5995-40 Fab fragments to bind simultaneously to a single protein. Importantly, this conserved *WxK* motif is present across multiple choline-binding proteins, providing a structural basis for the broad cross-reactivity observed. These findings explain both the extensive serotype breadth and the enhanced bacterial surface binding mediated by mAb 5995-40. Collectively, this work identified the conserved choline-binding region as a promising target for therapeutic and vaccine strategies aimed at providing improved protection against pneumococcal infection.

Overall, our study adds support for using human mAbs to highly conserved surface antigens for the prevention and treatment of pneumococcal infection. The application of human mAbs against *S. pneumoniae* also aids in the understanding of pneumococcal virulence factors and future immunogenic vaccine candidates for prevention of pneumococcal disease. Further defining protective epitopes of these mAbs will facilitate the development of a serotype-independent, broadly protective pneumococcal vaccine. Use of human mAbs for other bacterial infections is an important path forward for the development of new therapeutics across the field of bacteriology.

## Materials and Methods

**Key Resource Table**

| REAGENT or RESOURCE | SOURCE | IDENTIFIER |
| --- | --- | --- |
| **Antibodies** |  |  |
| APC anti-human CD19 | Biolegend | 302212<br>RRID: AB_314242 |
| Goat Anti-Human IgM-FITC | Southern Biotech | 2020-02<br>RRID: AB_2795600 |
| Goat Anti-Human IgD-FITC | Southern Biotech | 2030-02<br>RRID: AB_2795624 |
| Ghost Dye Red 710 | Cytek Biosciences | 13-0871-T100 |
| DAPI | Thermofisher | 62248 |
| PE Streptavidin | Biolegend | 405203 |
| Goat Anti-Human IgG Fc-AP | Southern Biotech | 2048-04 |
| Goat Anti-Human IgG HRP | Southern Biotech | 2048-05 |
| 5217-4, 5217-6, 5217-10, 5217-12, 5217-22, 5217-27, 5217-28, 5995-40, 5995-24, 5995-1702, 5995-2002, 5995-2116, 6100-42, 6100-61, 6100-294, 6100-888, 6100-1209, 6100-1311, 6100-1337, 6100-1505, 6100-1538, 6100-1873, 6100-1963 | This paper | N/A |
| PhtD3 | PMID: 33649050 | N/A |
| MPV481 | PMID: 35696580 | N/A |
| Goat Anti-human IgG (H+L) Cross-Adsorbed Secondary Antibody, Alexa Fluor 594 | Thermofisher | A-11014 |
| Goat Anti-Mouse C3 | Immunology Consultants Laboratory | GC3-90P-Z |
| **Bacterial and virus strains** |  |  |
| SPEC1 | BEI resources | Cat: NR-13388 |
| WU2 | Gift from Moon Nahm, University of Alabama Birmingham | N/A |
| TIGR4 | Gift from Larry McDaniel, University of Mississippi Medical Center | N/A |
| TIGR4ΔpcpA | Gift from Jorge Vidal, University of Mississippi Medical Center | N/A |
| TREP6A | BEI Resources | NR-51852 |
| SPEC6B | BEI Resources | NR-51853 |
| EMC9V | BEI Resources | NR-51855 |
| OREP10A | BEI Resources | NR-31703 |
| TREP11A | BEI Resources | NR-31705 |
| TREP12F | BEI Resources | NR-31704 |
| OREP18C | BEI Resources | NR-51856 |
| TCH8431 | BEI Resources | HM-145 |
| SPEC19F | BEI Resources | NR-51858 |
| EMC23F | BEI Resources | NR-51859 |
| ABC02110133 | CDC ABC Isolate Bank | N/A |
| A/California/07/2009 | IRR | Car: FR-201 |
| B/Washington/02/2019 | IRR | Cat: FR-1709 |
| RSV A2 | BEI Resources | Cat: NR-12149 |
| **Chemicals, peptides, and recombinant proteins** |  |  |
| PcpA | GenBank |  |
| PspA | GenBank |  |
| PsaA | GenBank |  |
| PhtE | GenBank |  |
| Ply | GenBank |  |
| NanA | GenBank |  |
| PcsB | GenBank |  |
| StkP | GenBank |  |
| PiuA | GenBank |  |
| LytA | GenBank |  |
| LytB | GenBank |  |
| PspC | GenBank |  |
| CbpD | GenBank |  |
| CbpE | GenBank |  |
| CbpJ | GenBank |  |
| Opti-MEM I + GlutaMAX | ThermoFisher | 51985034 |
| Freestyle 293 media | ThermoFisher | 12338026 |
| HiTrap Protein G column | Cytiva | 17040503 |
| HisTrap Excel column | Cytiva | 17371206 |
| HiTrap MabSelect column | Cytiva | 28408253 |
| PEI Max | Kyfora | 24765 |
| LB Broth | Thermofisher | AAH2667636 |
| LB Agar | Thermofisher | 611895000 |
| Kanamycin | Thermofisher | J61272.14 |
| Ampicillin | Thermofisher | J63807.09 |
| Streptomycin | Sigma | S-6501 |
| **Critical commercial assays** |  |  |
| ExpiFectamine CHO transfection kit | ThermoFisher | A29129 |
| E.Z.N.A Plasmid DNA Maxi Kit | Omega Bio-Tek | D6922-04 |
| Pierce Fab Preparation Kit | ThermoFisher | 44985 |
| **Deposited data** |  |  |
| Structure coordinates | PDB | 12QJ |
| CryoEM maps | EMDB | EMDB-76684<br>EMDB-76685 |
| Experimental models: Cell lines |  |  |
| Freestyle 293-F cells | ThermoFisher | R79007<br>RRID: CVCL_D603 |
| HL-60 cells | ATCC | CCL-240 |
| BEAS-2B cells | ATCC | CRL-3588 |
| Experimental models: Organisms/strains |  |  |
| C57BL/6 mice | Charles River<br>Laboratories | Strain Code: 027 |
| Software and algorithms |  |  |
| cryoSPARC | https://cryosparc.com/ | N/A |
| ChimeraX | https://www.rbvi.ucsf.edu/chimerax/ | N/A |
| AlphaFold3 | https://alphafoldserver.com/welcome | N/A |
| GraphPad Prism 10 | https://www.graphpad.com/ | N/A |
| Coot | https://bernhardcl.github.io/coot/ | N/A |
| Phenix | https://www.phenix-online.org/ | N/A |
| DeepEMhancer | https://github.com/rsanchezgarc/deepEMhancer | N/A |
| FlowJo | https://www.flowjo.com | N/A |
| Other |  |  |
| Superdex200 10/300 SEC column | Cytiva | 28990944 |
| Polyclonal serotype 4 serum | Statens Serum<br>Institute (SSI) | 16747 |
| International Standard for Human Anti-Pneumococcal Capsule Reference Serum | National Institute for<br>Biological Standards<br>and Control (NIBSC) | 007sp |
| Fibronectin, Human | Corning | 354008 |
| Collagen I, Bovine | Corning | 354231 |
| Bovine Serum Albumin Fraction V (BSA) | Roche | 03117332001 |
| Airway Epithelial Cell Basal Medium (AEBM) | ATCC | PCS-300-030 |
| Bronchial Epithelial Cell Growth Kit | ATCC | PCS-300-040 |
| Penicillin-Streptomycin-Amphotericin B Solution | ATCC | PCS-999-002 |
| Trypsin-EDTA Solution, 1X | ATCC | 30-2101 |
| Polyvinylpyrrolidone (PVP) | MP Biomedicals | 0219545180 |
| Iscove's Modified Dulbecco's Medium (IMDM) | ATCC | 30-2005 |
| Penicillin-Streptomycin solution (10,000 U/mL) | Gibco | 15-140-122 |
| Fetal Bovine Serum | Corning | MT35010CV |
| Dimethyl Sulfoxide (DMSO) | ATCC | ATCC 4X |
| Hanks' Balanced Salt Solution 10X (HBSS) | Gibco | 14065056 |
| Baby Rabbit Complement | Pel-Freez Biologicals | 31061-1 |

### Ethics Statement

All animal studies performed were in accordance with guidelines of the Institutional Animal Care and Use Committee of Florida State University under the Animal Use Protocol IPROTO202300000019, or the University of Georgia Animal Institutional Care Committee (protocol A2021 06-009 and A2023 11-002). All human subject blood isolations were performed at Emory University in accordance with the Institutional Review Board under the Protocol 2025P011087. Cone blood filters are from Shepeard Blood Bank in Augusta, GA and are not human subjects research.

### Blood draws and isolation of PBMCs

PBMCs from Subject 1 were obtained from leukocyte reduction filters supplied by Shepeard Community Blood Center in Augusta, Georgia as discarded samples. Each filter was gently washed with PBS (Corning, Cat# 21040CV) and the flow-through, which contained the leukocytes, was collected in a 50 mL conical tube. In a separate tube, 13 mL of warmed Ficoll-Paque (Cytiva, Cat #17144002) was added, and the cells were gently layered to the top of the Ficoll layer and centrifuged at 300 x g for 30 min at 4 °C with a slow acceleration and deceleration. The PBMC layer below the PBS and above the Ficoll was gently aspirated from the tube and added to a separate tube containing Dulbecco’s Modified Eagle’s Medium (DMEM) (Corning, Cat #10014CV) and centrifuged at 300 x g for 5 min. The supernatant was discarded, and the cell pellet was resuspended in 10 mL of red blood cell (RBC) lysis buffer (155 mM NH_4_Cl (Thermofisher, Cat #A661-500), 12 mM NaCO_3_, (Thermofisher Cat #S263-500) 0.1 mM EDTA (Thermofisher, Cat #BP2482100). The cells were centrifuged at 300 x g for 5 min, and the PBMCs were washed with DMEM. Finally, the cells were resuspended In ClonaCell-HY Medium A (StemCell Cat #03801) supplemented with 10% DMSO, transitioned to the -80 °C, and stored in the liquid nitrogen until further use. PBMCs from Subject 2 were obtained through study recruitment at Emory University Hospital after positive *S. pneumoniae* infection and written consent was obtained. Peripheral blood mononuclear cells (PBMCs) were isolated from human blood samples using the same Ficoll-Paque density gradient centrifugations as mentioned above, and PBMCs were frozen in the liquid nitrogen vapor phase until further use.

### Pneumococcal Protein Cloning and Expression

Protein sequences were retrieved from the genome of *S. pneumoniae* strains TCH8431 (serotype 19A) or TIGR4 (serotype 4) available in the NCBI database. The full-length proteins were cloned into the pET28a or pET29b vectors by Twist Biosciences (San Franscisco, CA). Once plasmids were obtained, each plasmid was transformed into *E. coli* BL21(DE3) (Thermofisher, Cat #EC0114) for protein expression. The following day, a single transformed colony was picked and cultured in 5 mL of LB broth (Thermofisher, Cat #AAH2667636) supplemented with antibiotic (50 µg/mL kanamycin for both pET28a and pET29b vectors) overnight in a shaking incubator at 37 °C. The overnight culture was then expanded in 250 mL of LB broth with antibiotic, cultured at 37 °C for 6 hrs, and then induced overnight with 50 μM isopropyl-d-thiogalactopyranoside (IPTG) (Thermofisher, Cat #BP1755-1) at 18 °C. Bacterial pellets were collected by centrifugation of 10,000 x g for 10 mins and resuspended in 50 mL of Lysis Buffer (50 mM Tris pH 8.0 (Corning, Cat #46-031-CM),150 mM NaCl (Thermofisher, Cat #AAJ21618A1), 0.5% TritonX-100 (Thermofisher, Cat #A16046AP), 5mM EDTA, 5mM DTT (Thermoscientific, Cat #FERR0861) and lysed by sonication. The cell lysates were centrifuged at 10,000 x g for 10 mins, and the supernatant was subsequently used for protein purification through a HisTrap column (Cytiva, Cat #17524801) following the manufacturer’s protocol.

Prior to antigen-specific sorting, 9 pneumococcal proteins from Table 1 (PcpA, PspA, PsaA PhtE, Ply, NanA, PcsB, StkP, PiuA) were biotinylated through the use of the EZ-Link NHS-PEG4 Biotinylation kit as per the manufacturer’s protocol (ThermoFisher, Cat #21455). Streptavidin-conjugated PE (Invitrogen, Cat #S866) was slowly added to the biotinylated antigens at a fluorophore to protein molar ratio of 4:1, adding 1 μL of streptavidin-PE every 20 minutes for a total of 5 additions and stored on ice away from light. PBMCs were suspended in FACS buffer (PBS, 2% FBS (Gibco, Cat #A5256701), 2% goat serum (Gibco, Cat #16210072), 0.5M EDTA) and Fc-blocked with Human TruStain FcX (BioLegend, Cat #422301) on ice for 30 min. A staining mixture containing anti-human CD19-APC (BioLegend, Cat #302212O), IgM-FITC (Southern Biotech, Cat #2020-02), IgD-FITC (Southern Biotech, Cat #2030-02), GhostDye Red 780 (Tonbo Biosciences, Cat #130865T100) or DAPI (ThermoFisher, Cat #62248) (live dead stains), and BT-SA conjugated antigen-PE was prepared and kept on ice away from light. Cells were centrifuged at 1100 x g for 5 min to pellet the cells and washed with FACS buffer once before resuspending in 30 µL FACS buffer. Single-stain controls were prepared with the cells and the remainder of the cells were added to the staining mixture and incubated on ice for 30 min away from light. Cells were washed once with FACS buffer and resuspended in PBS. Antigen-specific B cells (CD19 +, IgM-, IgD-, PE +, BV605 +) were single-sorted into a single tube and processed through 10X single cell sequencing following the manufacturers protocol.^48^ Single-sorted cells were processed using the Chromium GEM-X Single Cell 5’ Reagent Kits v3 on the Chromium X machine. After 10X processing and analysis, paired heavy and light chain plasmids were obtained and cloned by Twist Biosciences (San Francisco, CA) into their respective cloning vector (heavy: pTwist CMV hIgG1, lambda: pTwist CMV hIgL2, kappa: pTwist CMV hIgK). Once heavy and light chain plasmids were obtained, each plasmid was transformed into *E. coli* DH5a (Invitrogen, Cat #18258012) competent cells prior to plasmid isolation by maxiprep (Omega, Cat #D6922-04) per the manufacturer’s protocol. The antibodies were expressed in Expi293F cells (ThermoFisher, Cat #A14527) as previously described^49^. The culture supernatant was filtered through a 0.45 um filter (Thermofisher, Cat #FB12566511) before purifying through a HiTrap Protein G prepacked columns (Cytiva, Cat #17040401) per the manufacturer’s instructions. Purified antibodies were buffer exchanged into PBS and stored at -80 °C until further use.

### ELISA for Binding to Pneumococcal Proteins

To determine which recombinant antigen each mAb bound to, 384-well plates were coated with a pneumococcal antigen diluted to 2 µg/mL in PBS at 4 °C overnight. The plates were washed 1x with ddH_2_O and blocked with 2% blocking buffer (2% nonfat dry milk powder (ThermoScientific, Cat #NC9952266) dissolved in 0.05% PBS-Tween-20 with 2% goat serum added) for 1 hr at 37 °C. The plates were washed 3x with ddH_2_O and diluted mAb was added and incubated for 1 hr at 37 °C. The antibodies were diluted three-fold from an initial 20 µg/mL concentration in 1% blocking buffer (2% blocking buffer diluted 1:1 in PBS). The plates were washed 3x with ddH_2_O, and a goat anti-human IgG Fc-AP secondary antibody (Southern Biotech, Cat #1033-04), diluted 1:4000 in 1% blocking buffer, was added and incubated for 1 hr at 37 °C. The plates were washed 5x with 0.05% PBS-Tween-20. P-Nitrophenyl phosphate (PNPP) substrate (Thermofisher, Cat #34045) was added to substrate buffer (1.0M Tris base (Thermofisher, Cat #BP152-500), 0.5 mM MgCl_2_ (Thermofisher, Cat #M33-500) pH 9.8) to a concentration of 1 mg/mL and added to the plates. The substrate was incubated for 1 hr in the dark at room temperature, and then subsequently read at 405 nm on a BioTek plate reader.

### Biolayer Interferometry for Binding to Pneumococcal Proteins

To determine binding kinetics of mAb 5995-40 to each antigen, an initial baseline in running buffer was obtained for all biosensors. Following baseline, 100 µg/mL of His-tagged antigen was immobilized on anti-penta-HIS biosensor tips (Gator Biosciences, Cat #160009) for 120 sec. Then, the baseline signal was measured again for 60 sec before the biosensor tips were introduced into wells containing 100 µg/mL of primary antibody (5995-40) for 300 sec to determine the rate of association of mAb to antigen. To determine the rate of dissociation, the biosensors were introduced back into running buffer for 300 sec. Data collection and analysis were completed on the Gator Prime BLI machine (Gator Biosciences, Palo Alto, CA).

### Bacterial Strains and Growth Conditions

Pneumococcal strains were grown at 37 °C in 5% CO_2_ in Todd-Hewitt broth (Thermofisher, Cat #CM0189B) supplemented with 0.5% yeast extract (Thermofisher, Cat #H2676936P) for 16 hours. 20% glycerol (Thermofisher, Cat #G33-500) was added to the media, and 500 µL aliquots were made and stored at -80 °C until used. Prior to being used in experiments, the cultures were washed twice with 1 mL of PBS and resuspended in 500 µL of PBS. To determine the titer of bacteria, colonies were grown on BD Trypticase soy agar II with 5% sheep blood (Thermofisher, Cat #R060312). The cultures were serially diluted ten-fold, and the numbers of CFU/mL were determined. The calculated concentration was used for further experiments from aliquots thawed at later time points. After each experiment, the actual concentration of bacteria administered was determined through back-titration through plating on blood agar at the time of the assay. Strains used in this study are listed in Table 2.

### Bacterial Binding to Fixed Bacteria

The ability of the pneumococcal mAbs to bind to antigen exposed on the surface of *S. pneumoniae* was determined through flow cytometry. 1x10^6^ bacteria were incubated with 10 µg/mL of mAb for 30 min at 37°C. The bacteria were then washed with PBS plus 1% BSA (Thermofisher, Cat #BP9700100), and a secondary anti-human IgG Fc conjugated to Alexa Fluor 594 (Thermofisher, Cat #A-11014) at a 1:100 dilution incubated for 1 hour with the bacteria. Cells were washed with PBS-1% BSA and fixed in 2% paraformaldehyde (PFA) in PBS prior to analysis on the Cytek Aurora analyzer (Cytek Biosciences, Fremont, CA).

### Cell cultures of human respiratory cells

Human bronchial epithelial BEAS-2B cells (ATCC, Cat #CRL-3588) were cultured in tissue culture flasks pre-coated with a mixture of fibronectin (0.01 mg/mL) (Corning Cat #354008), bovine collagen type I (0.03 mg/mL) (Corning Cat #354231), and bovine serum albumin (BSA) (0.01 mg/mL) (Roche Cat #03117332001) dissolved in airway epithelial cell basal medium (AEBM) (ATCC Cat #PCS 300-030). Cells were maintained in complete growth medium consisting of airway epithelial cell basal medium (ATCC Cat #PCS 300-030) supplemented with a bronchial epithelial cell growth kit (ATCC Cat #PCS 300-040) , penicillin (10 U/mL), streptomycin (10 mg/mL), and amphotericin B (25 µg/mL) (ATCC Cat # PCS-999-002). Cells were incubated at 37°C with 5% CO_2_ and were supplemented with fresh medium 3 times weekly and passaged to a new flask once weekly or upon reaching ∼100% confluency by trypsinization using 0.25% trypsin–0.53 mM EDTA (ATCC 30-2101) solution containing 0.5% polyvinylpyrrolidone (PVP) (MP Biomedicals, Cat #0219545180) and subsequently seeded into experiment-specific devices. All experiments described hereafter were performed with cells that had been grown for ∼8-10 days in the specified device.

### Preparation of Inoculum

TIGR4, obtained from Dr. Jorge Vidal (University of Mississippi Medical Center, Jackson MS, USA) were recovered from frozen STGG stocks by streaking onto blood agar plates and incubating overnight at 37°C in 5% CO₂. Bacterial lawns were harvested in sterile phosphate-buffered saline (PBS; pH 7.4), and suspensions were adjusted to OD₆₀₀ = 0.1 (∼5.15 × 10⁸ CFU/mL) into the cell’s infection medium. Inoculum titers were verified by serial dilution and plating on blood agar plates. Unless specified otherwise, all *in vitro* experiments used freshly prepared inoculum.

### Infection of human respiratory cells with *S. pneumoniae* strains and treatment with mAb

Experiments were performed once the cell cultures became fully confluent and polarized, which occurred approximately 10 days post-seeding. Prior to infection, the pneumococcal inoculum was mixed with BEAS 2B infection medium and supplemented with 20 µg/mL of mAb 5995-40. Control bacterial inocula were not treated with the monoclonal antibody. Both treated and control preparations were incubated for 1 h at 37 °C in a 5% CO₂ atmosphere. BEAS 2B cells were washed three times with PBS, and the treated or control inoculum was then used to infect the cultures. Infected BEAS 2B cells were incubated for 1, 2, 3, or 4 h, after which medium from the basolateral compartment of the transwell device was collected, diluted, and plated onto blood agar plates to assess the burden of translocated pneumococci. Pneumococci attached to BEAS 2B cells were also harvested, and adhesion burden was determined by colony counts.

### HL-60 Cell Culture and Differentiation

The human promyelocytic leukemia cell line HL-60 (ATCC, Cat #CCL-240) was maintained in Iscove’s modified Dulbecco’s medium (IMDM; ATCC Cat. #30-2005) supplemented with 20% fetal bovine serum (FBS; Corning Cat. #MT35010CV), 100 U/mL penicillin-streptomycin (Gibco Cat. #15-140-122) at 37 °C in 5% CO_2_. Cells were cultured at 10^5^-10^6^ cells/mL. and differentiated into neutrophil-like cells using growth medium supplemented with 1.3% dimethyl sulfoxide (DMSO; ATCC Cat. #ATCC 4X) for 5-7 days. Prior to the opsonophagocytic killing assay (OPKA), HL-60 cells were washed, viability was assessed by trypan blue exclusion, and cells were resuspended at 2 × 10^5^ cells/mL in opsonization buffer (Hanks’ balanced salt solution [HBSS 10X; Gibco Cat. # 14065056] supplemented with 0.1% gelatin [Sigma; G6650] and 5% heat-inactivated FBS; hereafter referred to as OBB).

### Opsonophagocytic Killing Assay

The standard opsonophagocytic killing assay (OPKA) was performed according to the WHO pneumococcal antibody opsonization assay protocol (UAB-MOPA) provided by the WHO Pneumococcal Reference Laboratory,^50^ with modifications adapted for pneumococcal surface proteins.^51^ *S. pneumoniae* strains were grown to mid-log phase (OD₆₀₀ ≈ 0.2), washed, and adjusted to ∼1 × 10⁶ CFU/mL in OBB. Heat-inactivated polyclonal serum (56 °C, 30 min), or mAb 5995-40 were serially diluted (1:3) in OBB and incubated with the bacteria at 37 °C in 5% CO₂ for 30 min. Baby rabbit complement (Pel-Freez Biologicals; Cat. # 31061-1) either active or heat-inactivated for control wells, and differentiated HL-60 cells were then added, and plates were incubated for an additional 45 min at 37 °C in 5% CO₂ with shaking. Reactions were terminated on ice, and aliquots were plated onto THY agar. Colonies were enumerated after overnight incubation, and opsonophagocytic activity was calculated relative to complement control wells.

### Complement Deposition Assay

Antibody-dependent complement deposition was performed as previously described.^52^ In summary, biotinylated antigen was coupled with FluoSpheres NeutrAvidin beads (Invitrogen, Cat #F8776) at a 1:1 ratio of antigen (µg) to beads (µL). Beads were then washed in 5% PBS-BSA and resuspended at 1:10 of starting bead volume in 0.1% BSA. MAbs were then incubated at 1-10 µg/mL with 10 µL of antigen-specific beads for 2 hours at 37°C. Bead-mAb complexes were then washed twice in 0.1% BSA. Guinea pig complement (MP Biomedicals, Cat #8642836) was then diluted in R-10 buffer (RPMI-1640 with 10% FBS) at a 1:50 dilution. The complement and the bead-mAb complex were incubated for 15 minutes and washed twice in PBS. A 1:100 dilution of secondary anti-C3 antibody (ICL, Cat #GC3-90P-Z) was then incubated at room temperature for 15 minutes. Complexes were then washed twice in PBS and resuspended in a final volume of 150 µL PBS. The samples were read on a Cytek Aurora Analyzer.

### Pneumococcal Challenge

For the prophylactic intranasal challenge study, 4-6 week old C57BL/6 mice (Charles River Laboratory, Wilmington MA, strain #027) were used. Mice were intraperitoneally administered mAb treatments 2 hours prior to pneumococcal infection (n=10 mice per group). For infection, mice were anesthetized by inhalation of 5% isoflurane and intranasally challenged with either 1x10^7^ CFU of WU2 or 1x10^8^ CFU of TIGR4 in 40 µL of PBS. Mice were weighed and monitored for the presence of labored breathing and lethargy over the course of 14 days, with the humane euthanasia cutoff being 20% weight loss. For the therapeutic intranasal challenge study with WU2, 4-6 week old C57BL/6 mice (Charles River Laboratory, Wilmington MA, strain #027) were used. Mice were intraperitoneally administered mAb treatments 24 hours post pneumococcal infection (n=10 mice per group). For infection, mice were anesthetized by inhalation of 5% isoflurane and intranasally challenged with 1x10^7^ CFU of WU2 in 40 µL of PBS. To determine lung and blood bacterial titers, 4-6 week old C57B/6 mice (Charles River Laboratory) were intraperitoneally administered mAb 2 hours prior to pneumococcal infection and intranasally challenged with 1x10^7^ CFU of WU2 in 40 µL of PBS (n=10 mice per time point). At three different time points (24 hours, 48 hours, and 72 hours) post-infection, mice were euthanized, and blood and lungs were collected. Blood was collected via cardiac puncture and was serially diluted ten-fold and plated on TSA 5% sheep blood agar to determine bacterial titers. Lungs were extracted and homogenized in M tubes (Miltenyi Biotech, Cat #130-093-236) in 1 mL of PBS using the gentleMACs dissociator machine (Miltenyi Biotech, San Jose, CA). Lung homogenates were then serially diluted and plated to determine bacterial titers as above.

### Co-infection Challenge

For the IAV/SPN and IBV/SPN models, we utilized doses and timepoints as previously described^24^. 4-6 week old female C57BL/6 mice (Charles River Laboratory) were anesthetized by inhalation of 5% isoflurane and intranasally challenged with 100 PFU of H1N1 A/California/07/2009 or 100 PFU of B/Washington/02/2019 in 40 µL of PBS (n=15 mice per group). After 7 days, mice were anesthetized by inhalation of 5% isoflurane and intranasally challenged with 40 µL of 1x10^4^ CFU of pneumococcal strain WU2 in PBS. Two hours prior to bacterial challenge, mice were intraperitoneally administered 15 mg/kg of mAb 5995-40 or PBS. For the RSV/SPN model, we utilized doses and timepoints as previously described ^24^. 4-6 week old C57BL/6 mice were anesthetized by inhalation of 5% isoflurane and intranasally challenged with 2x10^6^ PFU of RSV A2 in 40 µL of PBS. After 5 days, mice were anesthetized by inhalation of 5% isoflurane and intranasally challenged with 40 µL of 1x10^6^ CFU of pneumococcal strain WU2 in PBS (n=20 mice per group). Two hours prior to bacterial challenge, mice were intraperitoneally administered 15 mg/kg of mAb 5995-40 or PBS. In all studies mentioned above, mice were weighed and monitored for the presence of labored breathing and lethargy over the course of 14 days, with the humane euthanasia cutoff being 30% weight loss (n=10 mice per group). Also, at two different time points (48 hours and 72 hours) post-infection, mice were euthanized, and blood and lungs were collected for all 3 co-infections (n=5 mice per group for IAV-pneumococcal and IBV-pneumococcal co-infections; n=10 mice per group for RSV-pneumococcal co-infections). Blood was collected via cardiac puncture and was serially diluted ten-fold and plated on TSA 5% sheep blood agar to determine bacterial titers. Lungs were extracted and homogenized in 1 mL of PBS using the gentleMACs dissociator machine (Miltenyi Biotech, San Jose, CA). Lung homogenates were then serially diluted and plated to determine bacterial titers as above.

### Size exclusion chromatography (SEC)

Individual CbpE protein, Fab, and Fab-antigen complexes were isolated through SEC on a Superdex S200 10/300 (Cytiva, Cat #28990944) in column buffer (120 mL NaCl, 20 mL Tris pH 7.5 (Corning, Cat #46-030-CM)) based on their molecular weight elution profiles. The desired eluent was concentrated and buffer exchanged into PBS before use.

### Assembly of Fab-antigen complex

5995-40 Fab was digested using the Pierce Fab Preparation Kit (Thermofisher, Cat #44985) prior to complexing with CbpE, according to the manufacturer’s protocol. Pure Fab and antigen were isolated through size exclusion chromatography (SEC) as described above. Fab and antigen were combined in a 2:1 molar ratio in PBS and incubated overnight at 4C. The Fab-antigen complex was isolated through SEC prior to use.

### Cryo-Electron Microscopy Data Collection

Cryo-EM grid preparation involved applying 4 µL of the sample to a Quantifoil 1.2/1.3 Cu 300 mesh grid, which had been plasma-cleaned for 20 seconds using a Solarus plasma cleaner (Gatan). Grids were plunge-frozen using a Vitrobot Mark IV (Thermo Fisher, Waltham, MA) at 10 °C and 100% humidity, with 1.5 s blot time and −1 blot force, then plunged into liquid ethane cooled by liquid nitrogen. After clipping and screening, data were collected on a Titan Krios equipped with a DE-Apollo direct electron detector. A single grid yielded 9,100 multi-frame micrographs. Images were recorded at a magnification of 59,000×, corresponding to a calibrated pixel size of 0.79 Å/pixel, with a total electron dose of 60 e⁻/Å². Another dataset with 30° angle tilt resulted 3,800 movies. All data was collected with Leginon.^53^

### Data Processing

Motion correction was performed using MotionCor3 for both tilted and unstilted movies. Corrected micrographs then imported into CryoSPARC to do the rest of cryo-EM data processing. Initial preprocessing included patch CTF estimation, followed by blob-based particle picking. After 3–4 rounds of 2D classification, well-resolved classes were selected, and three templates were generated for template-based particle picking. Particles were extracted with 2x binning (Fourier crop ½) and subjected to ab initio reconstruction to generate 5 distinct models. These served as references for multiple rounds (4–5) of heterogeneous refinement, progressively removing poorly aligned particles. At each step, only particles contributing to well-resolved classes were retained. The final particle stack was re-extracted without binning. One to two additional rounds of heterogeneous refinement were performed to further clean the dataset. The best particle sets from tilted and unstilted datasets then pooled before another round of heterogenous refinement to filter out the poorly aligned particles for the last time. The best ab initio model was used for non-uniform refinement. To mitigate orientation bias, the “Rebalance Orientation” job was used, followed by another round of non-uniform refinement with per-particle CTF refinement and defocus optimization enabled. Final maps were post-processed and sharpened using DeepEMhancer with the default “tight” model for figure preparation and visualization. Same procedure performed to group the particles into the class with multiple Fabs bound. The model with 3 Fabs showed the highest number of particles and highest resolution among others. We proceeded with these particles but not the model building due to the lack of enough resolution.

### Model Building and Refinement

Fab models were generated using AlphaFold3, and initial fitting of CbpE protein and Fab model into cryo-EM density maps was performed using UCSF Chimera’s *Fit in Map* tool.

Model refinement was conducted iteratively using PHENIX real-space refinement with manual adjustments in Coot. All refinements were performed against cryoSPARC-sharpened maps. The final models were evaluated for geometry and map fit using standard validation metrics.

### Data Deposition

Atomic coordinates were deposited in the Protein Data Bank (PDB) and the corresponding density maps in the Electron Microscopy Data Bank (EMDB). Motion-corrected micrographs were deposited in EMPIAR as the raw data files.

### Quantification and Statistical Analysis

One-way ANOVA was used to compare the complement deposition between the 5995-40 treated group and the negative control group (PIV04) using GraphPad Prism software (**GraphPad, San Diego, CA, USA**). A Welch’s t-test was used to compare the OPKA uptake between the 5995-40 treated group and the control serum groups. Two-way ANOVA was used to compare the bacterial translocation of TIGR4 in the 5995-40 treated group compared to the PBS treated group as well as the lung and blood bacterial titers for each pneumococcal infection and coinfection. Log-Rank (Mantel-Cox) was used to compare survival between all treated groups. Statistical significance was defined as follows: *, p ≤ 0.05, ** p ≤ 0.01; *** p ≤ 0.001; **** p ≤ 0.0001.

### Raw data

Raw data is available in FigShare at https://doi.org/10.6084/m9.figshare.32408832.

## Acknowledgements.

Influenza A Virus, A/California/07/2009, FR-201, was obtained through the International Reagent Resource, Influenza Division, WHO Collaborating Center for Surveillance, Epidemiology and Control of Influenza, Centers for Disease Control and Prevention, Atlanta, GA, USA. Influenza B Virus, B/Washington/02/2019 (Victoria Lineage), FR-1709, was obtained through the International Reagent Resource, Influenza Division, WHO Collaborating Center for Surveillance, Epidemiology and Control of Influenza, Centers for Disease Control and Prevention, Atlanta, GA, USA. *Streptococcus pneumoniae* multi-drug resistant serotype 3, ABC2020110133, were provided by the Active Bacterial Core surveillance (ABCs) Isolate Bank, a collaboration between the Centers for Disease Control and Prevention and state health departments through the Emerging Infections Program.

