## Supplementary Information for "A pan-serotype human monoclonal antibody protects against pneumococcal infection by targeting multiple choline binding domain proteins"

| Table S1. Cryo-EM data collection, refinement and validation statistics. | | |
|---|---|---|
| **Data collection and processing** | #1 one Fab bound | #2 three Fabs bound |
| Scope | Titan Krios | Titan Krios |
| Voltage (kV) | 300 | 300 |
| Electron exposure (e–/Å$^2$) | 60 | 60 |
| Defocus range (μm) | 0.8-2.8 | 0.8-2.8 |
| Camera | DE-Apollo | DE-Apollo |
| Pixel size (Å) | 0.79 | 0.79 |
| Symmetry imposed | C1 | C1 |
| Initial particle images (no.) | 1,100,000 | 1,100,000 |
| Final particle images (no.) | 58,000 | 93,800 |
| Map resolution (Å)<br>FSC threshold | 3.35<br>0.143 | 3.7<br>0.143 |
| Map resolution range (Å) | 3-6 | 3.5-8 |
| **Refinement** | | |
| Model resolution (Å)<br>FSC threshold | 3.35<br>0.143 | NA<br>0.143 |
| Model resolution range (Å) | 3-6 | NA |
| Map sharpening *B* factor (Å$^2$) | 111.3 | 158.6 |
| R.m.s. deviations<br>Bond lengths (Å)<br>Bond angles (°) | <br>0.004<br>0.839 | <br>NA<br>NA |
| Validation<br>Clashscore<br>Poor rotamers (%) | <br>21.22<br>4.79 | <br>NA<br>NA |
| Ramachandran plot<br>Favored (%)<br>Allowed (%)<br>Disallowed (%) | <br>83.17<br>16.03<br>0.80 | <br>NA<br>NA<br> |
| EMDB<br>PDB<br>EMPIAR | EMDB-76684<br>12QJ<br>TBD | EMDB-76685<br>NA<br>TBD |

| Table S2. Monoclonal Antibody Sequences. | | | |
|---|---|---|---|
| **mAb Name** | **Heavy Chain Sequence** | **Light Chain Sequence** | **Light Chain Type** |
| **5217-4** | gaggtgcagctggtggagtctgggggaggc<br>ttggtccagcctggagggtccctgagactctc<br>ctgtgcagtgtctggatttaccttcagtgacca<br>ctacatggactgggtccgccaggctccagg<br>gaagggactggagtggcttggccgtcttaga<br>cccagatctaataattacaacgcagattacg<br>ccgcgtctgtgaaaggcagattcaccatctc<br>aagagatgattcaaagaactcactgtatctg<br>caaatgaacagcctgaaaaccgacgacac<br>ggccatatattactgtgctagactcgcgggtt<br>gggttggacatgctgatgattttgatatatggg<br>gccaagggacactggtcaccgtctcttca | gacatccagatgacccagtctccatctt<br>ccctgtctgcatctgtaggagacagagt<br>caccatcacttgccgggcaagtcagg<br>acattagaaatgatttaggctggtatca<br>acaggaaccagggaaggcccctaag<br>cgcctggtctatgctgcatcgactttgca<br>aagtggggtcccatcaaggttcagcgg<br>cagtggatctgggacacaattcacgct<br>cacaatcagcagcctgcagcctgaag<br>attttgcaacttattactgtctgcagcatta<br>tcgttacccgtgcaccttcggccaagg<br>gaccaaggtggaaatcaaa | Kappa |
| **5217-6** | caggtgcagctggtggagtcgggggggaaa<br>cttggtcaagcctggcgggtccctgagactct<br>cctgtgcagcctctggattcacctttagcgact<br>tctacatgaattggattcgccaggctccagg<br>gaaggggctggagtgggtcgcacatgtgag<br>tggtcctggcaccaccaaatggtacgcagat<br>tctgtgatgggccgattcaccatatccaggg<br>acaacaccaagaattcactgcatctgcaaat<br>ggacagtcttacagccgaagacacggccct<br>atattactgttcgagagggatattttcggactc<br>ctggggccagggaaccctggtcaccgtctc<br>ctca | gacatcgtgatgacacagtctccagac<br>tccctgtctgtgtctctgggcgagaggg<br>cctccatcaagtgcaagtccagccag<br>actattttggacagttccaacaataaga<br>actttttagcttggtaccagcagaaacc<br>aggacagtctcctaagttactcttttactg<br>ggcatctacccgggaatctggggtccc<br>tgaccgattcagtggcagcgggtctgg<br>gacagatttcactctcaccatcagcag<br>cctgcaggctgaagatgtggcaatttat<br>ttctgccaccaatattatagtcttccgtgg<br>acgttcggccaagggaccaaggtgga<br>aatcaaa | Kappa |
| **5217-10** | caggtgcggctgcaggagtcgggcccagg<br>gctggtgaagccttcggagacactatccctc<br>acgtgcaatgtctctggtggctccgtcgacaa<br>tggacgttattactggagctggatccggctgc<br>ccccagggagacagctggagtggattggct<br>atttctatttcagtgggagcaccaaatacaac<br>ccctccctcgagagtcgtctctccatttcaggg<br>gacacagtcagcaaccaaatcttcctgcgc<br>ctcagttctgtgacagttgatgacacggccgt<br>ctactattgtgcgagtgaagttttacattgggg<br>ctttgaaagttggggccagggaaccctggtc<br>accgtctcctca | gacatccagatgacgcagtctccctcg<br>tccctgtctgcgtctgttggagacagagt<br>caccatcacttgccgggcaggagaga<br>acattagggagtatttgaattggtatcag<br>aaaaaacctgggagagcccctacact<br>cctgatctatgctgcagatagtttgcaag<br>gtggagtcccgccaaggttcagtggcg<br>ccggatctgggacagagttcaccctca<br>ccatcagcagcctgcaacctgaagatt<br>ttgcgacttattactgtcaacagagttac<br>aaaatccctcggacgttcggccaagg<br>gacaagggtcgaaatcaaa | Kappa |
| **5217-12** | caggttcaactgcaggagtcgggcccagg<br>actgatgaagccttcggggaccctgtccctc<br>acctgcggtgtctctggtgtctccatcagtagt<br>gataactggtggagttgggtccgccagtccc<br>cagataagggcctggagtggatcggggag<br>aggtatcatagtgggagcagcaactacaac<br>ccgtccctcaagagccgagtcaccatcaca<br>gtagacaagtccacgaaccagttctccctga<br>tgctgagctctgtgaccgccgcggacacgg<br>ccgtctattactgtgcgattttaaaccgtggga<br>cgtcgtggtactttgaacactggggccaggg<br>agccctggtcaccgtctcctca | cagtctgccctgactcagcctgcctccg<br>tgtctgggtctcctggacagtcgatcac<br>catctcctgcactggaaccagccgtga<br>tgttgggacttataaccttgtctcctggta<br>ccaacaacacccgggcaaagccccc<br>aaacttatgatttttgaggtcaataagcg<br>gccctcagcggtttctaatcgcttctctg<br>gctccaagtctggcaacacggcctccc<br>tgacaatctctgggctccaggctgagg<br>acgaggctgattattactgctgttcatat<br>gcaggtagtgccactttggtattcggcg<br>gagggaccaagctgaccgtcgta | Lambda |
| **5217-22** | caggtgcagctggtgcagtctggggctgag<br>gttaaggagcctggggcctcagtgaaggtct | cagtctgccctgactcagcctccctccg<br>cgtccgggtctcctggacagtcagtca | Lambda |

| | | | |
|---|---|---|---|
| | cctgcaaggcttctggatacaccttcagcgactactatatacactgggtgcgacaggcccctggacaggggcttgagtggatgggatggatcaaccctaacactggtggcacagactatgcacagaagtttcagggctgggtcaccatgaccagggacacgtccgtcagcacagcctacatggacttcagcgggctgaagtctgacgacacggccatatattactgtgcgaggtccggtatgatatcacgatattttgaccccagttggttcgaccccctggggccagggaaccctggtcaccgtctcctca | ccatctcctgcaccggagccagcactgacattggtaattataactatgtctcctggtaccaacaacacccaggcaaagccccaagctcattatttatgaggtcattaagcggccctcaggggtccctgatcgcttctctggcttcaagtatggcaacacggcctccctgatcgtctctgggctccaggctgaggatgaggctgattattactgcagctcatatgcaggcaagaacacttgggtgttcggcggagggaccaaactgaccgtccta | |
| **5217-27** | caggtgcagctggtggcgtctgggggaggcgtggtccagcctgggaggtccctgagactctcctgtgcagcctctggattcacatttaattcttatgctatgcattgggtccgccaggctccaggcaaggggccggagtgggtggcagttatttcatataacgcaaacaatgaatactacgcagactccgtgaagggccgattcaccatctccagagacaattccaaggacacactggatctgcaaatgaacagcctgacgactgaggacacggctgtgtattactgtgtgaaagatggccctcccagagtgagttctacggtgactacgggctggttcgaccccctggggccagggaaccctggtcaccgtctcctca | gacatccagatgacccagtctccatcttccctgtctgcatcagtggaagacagagtcaccatcacttgccgggcaagtcagaacattaaatattatttgaattggtatcagcagcaaccagggaaagcccctaaactcctgatctattctgcattcagtctgcaaagtggggtcccttcaaggttcagtggcagtggatctgggacagatttcactctcaccatcaccagtctgcagcctgaagattctgcaacctattactgtcaacagagtcacagtatccctttcacttttggccaggggaccaagcttgagatcaaa | Kappa |
| **5217-28** | gaggtgcagctggtggagtctgggggaggcttggtacagcctggagggtccctgagactctcctgtgcagcctctggattcaccttcagtagttatgaaatgaactgggtccgccaggctccagggaaggggctggagtgggtttcatacattagtagtagtggtagtaccatatactacgcagactctgtgaagggccgattcaccatctccagagacaacgccaagaactcactgtatctgcaaatgaacagcctgagagccgaggacacggctgtttattactgtgcgagttgggactacggtgactacgcccgatattactactactactactactacatggacgtctggggccaagggaccacggtcaccgtctcctca | gaaatagtgatgacgcagtctccagccaccctgtctgtgtctccaggggaaagagccaccctctcctgcagggccagtcagagtgttagcagcaacttagcctggtaccagcagaaacctggccaggctcccaggctcctcatctatggtgcatccaccagggccactggtatcccagccaggttcagtggcagtgggtctgggacagagttcactctcaccatcagcagcctgcagtctgaagattttgcagtttattactgtcagcagtataataactggcctccgcggactttcggccctgggaccaaagtggaaatcaaa | Kappa |
| **5994-40** | caggtgcagctgcaggagtcgggcccacgactggtaaaggtttcggagaccctgtcactcacctgctctctttctggtcactccatcagtagtggtcaccactgggcctggatccggcaatccccgggaggggattggagtggattgcgactgtcgatcataacgggaggacctactacaatccgtctctcaggggtcgcgtcatactgtcgaaagacacaaccaagaattccttctccgtgcagttgacttctgtgaccgccgcagacacggccgtctactactgtgcgagacgagaggccccttcacctggaactggctggtatcctccctttggacagtggggccagggagcgctggtcacggtctccaca | cagtctgtgctgacgcagccgccctcactctctggggccccagggcagacggtcaccatctcctgcactgggagcagctccaacatcggggcaggttatgatgtacactggtaccgacaacttccaggaacagcccccacactcctcatctctgtctttgataatcggcccttaggggtccctggccgaatctctggctccaagtctggcacctcagcctccctggccatcactgggctccaggctgaggatgaggctgactattactgccagtcccatgacaacatgttgagtggttcggtgttcggcggagggaccaagctgaccgtcctt | Lambda |
| **5995-247** | caggtgcaactgcaggagtcgggcccaggactggttaagccttcggagaccctgtccctcacctgcggtgtctctggttactacatcagtagtgctttctattggggctggatccggcagccccca ggaaaggggctggagtggattgccagtattgattccagcgggactacgtatcacaaccctc | cagtctgccctgactcagcctccctccgtgtccgcgtctcctggacagtcagtcaccatctcctgcactggaaccagcagtgacgttggtgcttatgaccgtgtctcctggtatcagcagcccccccggcacagtccccaaactcattatttttgaggtcactaatcggc | Lambda |

| | | | |
|---|---|---|---|
| | ctttgaggggtcgagtctccatgtcagtcgac<br>acgtccaggaaccagttttccctgaggctga<br>actctgtgaccgccgcagacacggccgtct<br>atttctgcgtgagaggggctccccgatattgtt<br>ctttctggggccagggaattctggtcagcgtc<br>tcctca | cctcaggggtccccggtcgcttctctgg<br>gtccaagtctgccaacacggcctccct<br>gaccatctctggactccggggtgagga<br>cgaggctgattattactgcagctcatatg<br>caaggggtggcattttgattttcggcgg<br>agggaccaggctgaccgtccta | |
| **5995-1702** | cagctgcagcttctgcagtcaggggctgag<br>gtgaagaagcctgggtcctcggtgaagattt<br>cttgcagggcttttgggggcaatttcgagagtt<br>atgccattgcctgggtgcgacaggcccctgg<br>acaaggccttgagtggatgggaggcattgtc<br>cccatcctctctgaaacaaagtacgcacag<br>aatttccaggacagactcaccattaccgcgg<br>accaagccacgagtacagtttacttgcaatt<br>gaacagcgtcacgtctgcggacacggccttt<br>tattactgtacgagagatgggcagtggctac<br>ggtctgacctgcagtatgaggctttcgatgtct<br>ggggccaggggacactggtcaccgtctcct<br>ca | cagtctgccctgactcagcctgcctccg<br>tgtctgggtctcctggacagtcgatcac<br>catctcgtgcactgggaccgacagtga<br>tcttgggagcgataaccgtgtctcctggt<br>accaacaacaccccggcacagcccc<br>caaactcatactttttgagatcactaaga<br>ggccctcaggagtttcttctcgcttctctg<br>gctccaagtctggcaacacggccttcct<br>gacaatctctgggctccaggctgggga<br>cgaggctgacttttactgctgctcatatg<br>caggtgacaatacttatgtcttcgggact<br>gggacccaggtctccgtccta | Lambda |
| **5995-2002** | gaggtgcagttgttggagtctgggggaggct<br>tggtacagcctggggggtccctgagactctc<br>ctgtgcagcctctggattcagctttagcagtta<br>tgccatgagctgggtccgccagggtccagg<br>ggaggggctggagtgggtctcttctattagtg<br>ctagcggtcgtagcacatactacgcagactc<br>cgtgaagggccgattcaccatctccagaga<br>caattccaggaatacgttgtatgtgctaataa<br>acagtctgagagccgaggacacggccgtct<br>atttctgtgcgaaatctcaccctcatagtagtg<br>gcgggtatggcgaaaactccaacctgtttga<br>ctactggggccagggaaccctggtcaccgt<br>ctcctca | gaagtcgtgctgacgcagtctccagcc<br>accctttctgtgtctccaggggaaagag<br>ccaccctctcctgcagggccagtcaga<br>gtgttagaagcaacttagcctggtaccg<br>gcagaaacctggccaggctcccagg<br>ctcgtcatctatggtgcatccaccaggg<br>ccactggtatcccagccaggttcagtg<br>gcagtgggtctgggacagagttcactct<br>caccatcagcagcttggagtctgaaga<br>ttttgcagtttattactgccagcagtatatt<br>aactggcctccagtcactttcggccctg<br>ggaccagagtggaaatcaaa | Kappa |
| **5995-2116** | gaggtgcagcttgtggagtctgggggaggct<br>tgatacagcctggggggtccctgagactctc<br>ctgtgcagcctctggattcaccttcagtagcta<br>tagcatgaactgggtccgccaggctccggg<br>gaagggcctggagtgggtttcatacatcagt<br>cctactagtactaccatttactacgcagagtct<br>gtgaagggccgattcaccatctccagagac<br>aatgccgagaactcactgtttctgcaaatga<br>acagcctgagagccgaggacacggctgtct<br>actactgtgcgagagatctgacggaggcat<br>cacggtggttcggggagctcccgaagggtg<br>ctcttgatatctggggccaagggacaacggt<br>caccgtctcttca | gaaattgtggtgacgcagtctccaggc<br>accctgtctttgtctccgggggaaagag<br>ccaccctctcctgcagggtcagtcaga<br>gtgttagcagcacttacttagcctggtac<br>cagcataaacctggccaggctcccag<br>gctcctcatctatggtgtatccaacagg<br>gccactggcatcccagacaggttcact<br>ggcagtgggtctgggacagacttcact<br>ctcaccatcagcagactggagcctga<br>agattttgcagtgtattactgtcagcagt<br>atcgtagttcaccgctcactttcggcgg<br>agggaccaaggtggagatcaaa | Kappa |
| **6100-42** | caggtgcaactggtgcagtcggggggagg<br>cctgggacacccatggggtccctgcgact<br>ctcctgtgcagccacgggattcgacttcaga<br>aactatgccatgacttgggtccgccagactc<br>cagggaggggactggagtgggtctcaagta<br>ttagtggcaacgatgagtggacatattatgca<br>gactccgtgaagggccgcttcaccgtctcca<br>gagacaacgccaggaacatctttttattgcat<br>ctggacagcctgagagtcggggacacggc<br>cctttatttttgtgcgagaggggaaccgcgag | gacatccagatgacccagtctccatcct<br>cactgtctgcatttgttggagacaaagtc<br>accctcacttgtcgggcgagtcaggac<br>attgaaaattatctggtctggtttcagca<br>gcaaccagggaaacccccgaagtcc<br>ctgatctatgctgcatcccgtctacaga<br>gtggagtgtcgtccaggttcagcggca<br>gcggatctgggacacatttcactctcac<br>catcagcaggctgcagcctgcagattt<br>cgcaacatattattgtcaacaatatcac | Kappa |

| | | | |
|---|---|---|---|
| | gtacctatgtcttcaattactggggccagggagtcctggtcaccgtctcctca | agtttccctccgaccttcggcccggggacacgactggagattaaa | |
| **6100-61** | caggtggagctactacagtggggcgcaggactgttgaggccctcggagaccctgtccctcacctgcgttgtctctgatgggtccttcactgcttactactggagctggatccgccaggccccagggaaggagccggagtggattggggaagtccatcataatagacacaccatctacaacccgtccctcgagagtcgagttacgatattatcagactcgtccagcagccaattctccctgatgctgacctctgtgaccgccgcggacacggctgtatactactgtgcgagacactatggacgtactcttgactactgggtacagggaaccctggtcaccgtctcctca | aatattgtgatgacccaggctccactctcctcacctgtcacccttgggcagccggcctccatctcctgcaggtctaatcaaagcctcgtacacagtgatggatacacctacttgagttggtttcaccagaggccaggccagcctccaagactcctaatttatgcgctttctaaccgcttctctggggtcccagacagattcagtggcagtggggcagtgacagatttcacactgagaatcagcagggtggaagctgaggatgtcgggatttattactgcatgcaatctacacaatttcctcggacctttggccaggggaccaagctggagatcaaa | Kappa |
| **6100-294** | caggtccagctggtacagtccggggctgaggtgaagaagcctggggccaccgtgaaaatctcctgcaaggcttctggatacaccttcaccgactactacatgcactgggtccaacaggcccctggaaaagggcttgagtggatgggatatgttgatcctgaagatggtgaaacaatctctgcagagaagttccagggcagagtcaccataaccgcagacacgtctacagacacagtctacatggagctggccagcctgagatctgaggacacggccgtgtattactgtgccccgggggattacgatgcccttatctggggccaagggacaatggtcaccgtctcttca | cagcttgtgctgactcaatcgccctctgcctctgcctccctgggagcctcggtcaagctcacctgcactctgagcagtgggcacagcagctaccccatcgcatggcatcagcagtcgccagagaagggccctcggtttttgatgaggcttaacagtggtggcgatcacaccaaggggacgggatccctgatcgcttctcaggctccagctctggggctgagcgctacctcaccatctccagcctccagtctgaagatgaggctgactattactgtcagacctggggcactggcacctgggtgttcggcggagggaccaagctgaccgtccta | Lambda |
| **6100-888** | cagctggtacagtctggggctgaggtgaagaaacctgggtcctcggtgaaggtctcctgcaagacttctggagacaccttcagcaatcacccctatcagctgggtgcgacaggcgcccggacacgggcttgagtggatgggcaggatcttctctctctttggtacaacggagtacactcagaactttcaggacagaatcgcccttaccgcggacgaatctacgaccacagtctacatggaactgagcagcctgagatctgacgacacggccgtgtattactgtgcgagagagggcagtggctcgagggtccatgcttttgatatctggggccaggggactatggtcaccgtctcatca | gacatccagatgacccagtctccttccaccctgtctgcatctgtaggagacagagtcatcatcacttgccgggccagtcagagtattagtacctggttggcctggtatcagcagaaaccagggaaagcccccaagctcctgatctatcaggcgactaatttagaaagtggggtcccgtcaacgttcagcggcagtggatctgggacagaattcactctcaccatcagcagcctgcagcctgatgattttgcaacttattactgccaacaatataaaacttataggacgttcggccaagggaccaaggtggaaatcaaa | Kappa |
| **6100-1209** | caggtgcagttggtgcagtctggggctgaggtgaagaagcctggggcctcagtgaaggtctcctgcaaggcttctggatacacgttcgccggctactatctgcattgggtgcgacaggcccctggacaagggcttgagtggatgggacgggtcaatcccaccagcggtggcatcaagtatgcacagaactttcagggcagggtcaccatgacccgcgacacgtccaccggcacagcctacatggagttgaagaggctgacatctgacgacacggccatatatttctgtgcgcgagagggggaggaaacaccgacggtcccctactttgactactgggccagggaacactggtcaccgtctcctca | cagcctgtgctgactcagccaccttcctcctccgcatctcctggagaatccgccagactcacctgcaccttgcccagtgacctcaatgttggtgggtacaacatttactggtatcagcagaagccagggagccctcccaggtatctcctgttttactactctgactcaagtaagggccagggctctggagtccccagccgcttctctggatccaaagatgcttcagccaatgcagggattttgctcatatccgggctccagtctgaggatgaggctgactattactgtatgatttggccaaggaatgcgcctgtggtattcggcggagggaccaagctgaccgtccta | Lambda |
| **6100-1311** | caagtgcagctgatggagtctgggggaggcttggtacagcctggcaggtccctgagactctcctgtgcagcctctggattcacatttgatgattat | gacatccagatgacccagtctccttccaccctgtctgcatctgttggagacagagtcaccatcacttgccgggccagtcaga | Kappa |

| | | | |
|---|---|---|---|
| | gccatgcactgggtccggctagttccaggga<br>agggcctggagtgggtctctgggattaattgg<br>aatagtgataccatagactatgcggactctgt<br>gaagggccgattcaccatctccagagacaa<br>cgccaagaactccctatatctgcaaatgagc<br>agtctgagagctgaggactcggccttgtatta<br>ctgtgcaaaggatattgcagctcgtccttact<br>actactactacatggacgtctggggcaaag<br>ggaccacggtcaccgtctcctca | gtattagttcctggttggcctggtatcagc<br>agaaaccagggaaagcccctgagct<br>cctgatctatcaggcatctagtttacaaa<br>ctggggtcccatcaaggttcagcggca<br>gtggatctggaacagaattcactctcgc<br>catcagcagcctgcagcctgatgatttt<br>gcaacttattactgccaacaatatagtc<br>attcctcgtggacgttcggccaaggga<br>ccagggtggaaatcaaa | |
| **6100-1337** | caggtgcagctggtggagtctgggggaggc<br>ttggtcaagcctggagggtccctgagactctc<br>ctgtgcagcctctggattcaccttcagtgacta<br>ctccatgagctggatccgccagcctccagg<br>gaggggactggagtgggtttcatacattacta<br>gtgatgataataccatatactatgcagactttg<br>tgaagggccgattcaccatctccagggaca<br>acgcccagaactcactgtatctgcaagtgaa<br>cagcctgagagccgaggacacggccgtat<br>attactgtgcgagagatcttctgcctggttatc<br>gcaccgctgaatacttccaacactggggcc<br>agggcaccctggtcaccgtctcctca | tcctatgtgctgactcagccaccctcagt<br>gtcagtggccccaggacagacggcc<br>aggattacctgtgggggaaacaacatt<br>ggaagtaaacctgttcactggtatcagc<br>agaggccaggccaggcccctgtgctg<br>gtcgtctttgatgatagcgaccggccct<br>cagggatccctgagcgattctctggctc<br>caactctgggaacacggccaccctga<br>tcatcaccagggtcgaggccggggat<br>gaggccgactattactgtcaggtatgg<br>gatattagtagtgatcatccgaaggtga<br>tattcggcggagggaccaggctgacc<br>gtccta | Lambda |
| **6100-1873** | caggtgcagctggtggagtctgggggaggc<br>ctggtcaagcctggggggtccctgagactct<br>cctgtgcagcctctggattcagcttcagttact<br>atagcatgaagtgggtccgccaggctccag<br>ggaaggggctggaatgggtctcatccattatt<br>actagtagtgattacatatactacgcagactc<br>agtgaagggccgattcaccatctccagaga<br>caacgccaagaactctctgtatctgcaaatg<br>aacagcctgagagccgaggatacggctgt<br>gtattactgtgcgagagctgccggacattgta<br>gtggtgcagtgggggactactgctactacat<br>ggacgtctggggcaaagggaccacggtca<br>ccgtctccagc | cagtctgtgttgacgcagccgccctca<br>gtgtctgcggccccaggacagaaggt<br>caccatctcctgctctggaagcagatcc<br>aacattgggaaaaattttgtttcctggta<br>ccagcaactcccaggaacagccccc<br>aaactcctcatttatgactataataagcg<br>accctcagggattcctgaccgattctct<br>ggctccaagtctggcacgtcagccacc<br>ctgggcatcaccggactccagactgg<br>agacgaggccgattattactgcggaac<br>atgggatatcagcctgagtgcctgggt<br>gttcggcggagggaccaagctgaccg<br>tccta | Lambda |
| **6100-1953** | caggtgcagcttgtggagtctgggggaggct<br>tgatacagcctggggggtccctgagactctc<br>ctgtgcagcctctggattcaccttcaatggcta<br>tagcatgaactgggtccgccaggctccggg<br>gaagggcctggagtggatttcatacattagtc<br>ctactagtactaccatttactacgcagagtctg<br>tgaagggccgattcaccatctccagagaca<br>atgccgagaactcactgtttctgcaaatgaa<br>cagcctgagagccgaggacacggctgtcta<br>ctactgtgcgagagatctgacggaggcatc<br>acggtggttcggggagctcccgaggggtgc<br>tcttgatatctggggccaagggacaacggtc<br>accgtctcttca | gacattgtggtgacgcagtctccaggc<br>accctgtctttgtctccgggggaaagag<br>ccaccctctcctgcagggtcagtcaga<br>gtgttagcagcacttacttagcctggtac<br>cagcataaacctggccaggctcccag<br>gctcctcatctatggtgtgtccaacagg<br>gccactggcatcccagacaggttcact<br>ggcagtgggtctgggacagacttcact<br>ctcaccatcagcagactggagcctga<br>agattttgcagtgtattactgtcagcagt<br>atcgtagttcaccgctcactttcggcgg<br>agggaccaaggtggagatcaaa | Kappa |

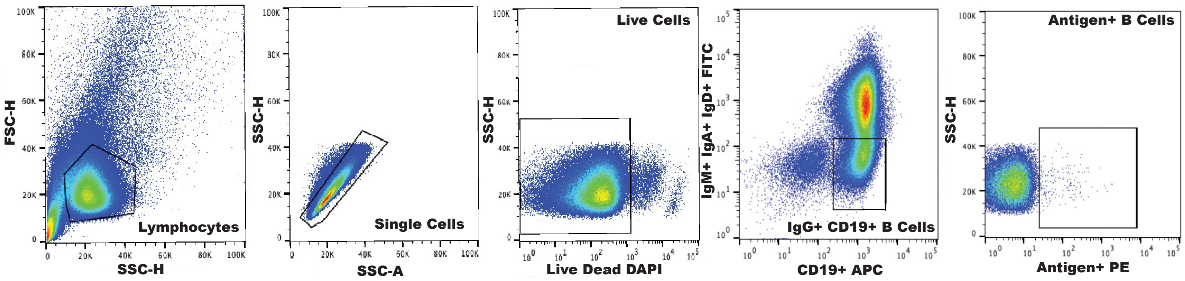


**Figure S1. Flow Panel to Isolate Antigen-specific B cells**. PBMCs were sorted according to the panel above to obtain live CD19$^{+}$/IgD$^{-}$/IgM$^{-}$/IgA$^{-}$ antigen-specific B cells.

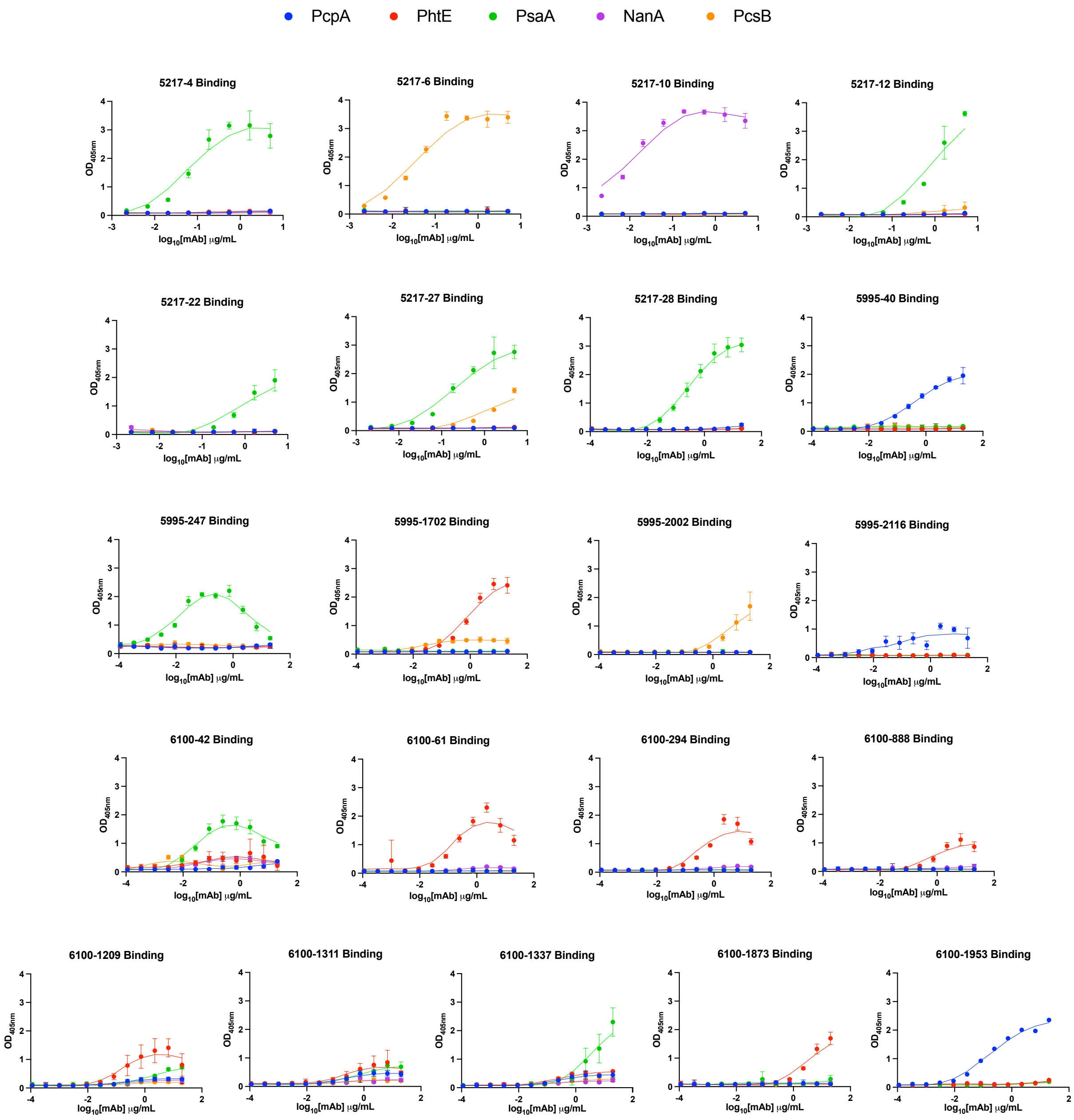


**Figure S2: ELISA curves for each tested antibody.** Recombinantly expressed monoclonal antibodies (mAbs) were tested against nine antigens, of which only 5 antigens demonstrated measurable mAb binding. Panels show the individual binding curves for each mAb against the 5 antigens that demonstrated antibody-antigen binding with each point on the graph correlating to the $OD_{450}$ measured at that given mAb dilution. Error bars are the standard deviation of four technical replicates at each mAb dilution.

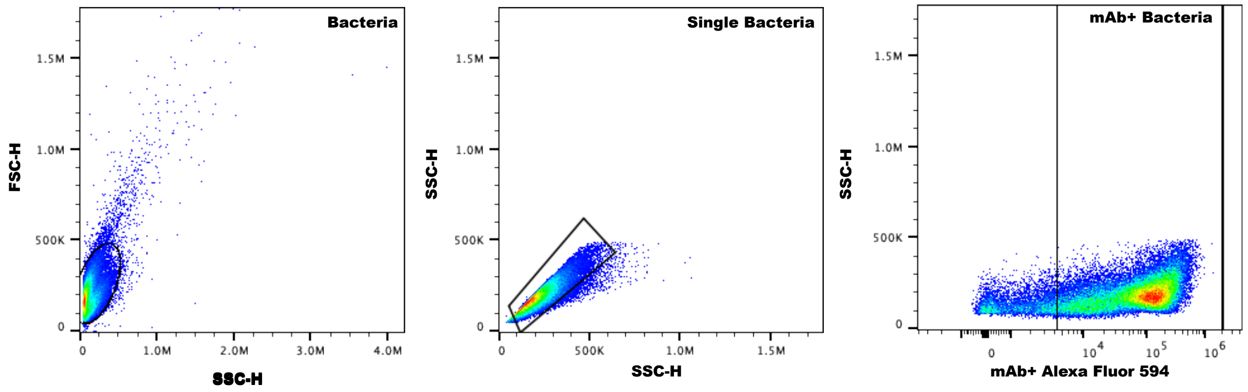


**Figure S3: Bacterial binding example flow panel.** Multiple *S. pneumoniae* serotypes were used to assess binding of mAb 5995-40 to the bacterial surface. Following incubation with mAb 5995-40, an Alexa Fluor 594 (AF594)-conjugated anti-human Fc secondary antibody was added. Samples were analyzed by flow cytometry using the gating strategy shown above, and binding was quantified as percentage of AF594+ bacteria.

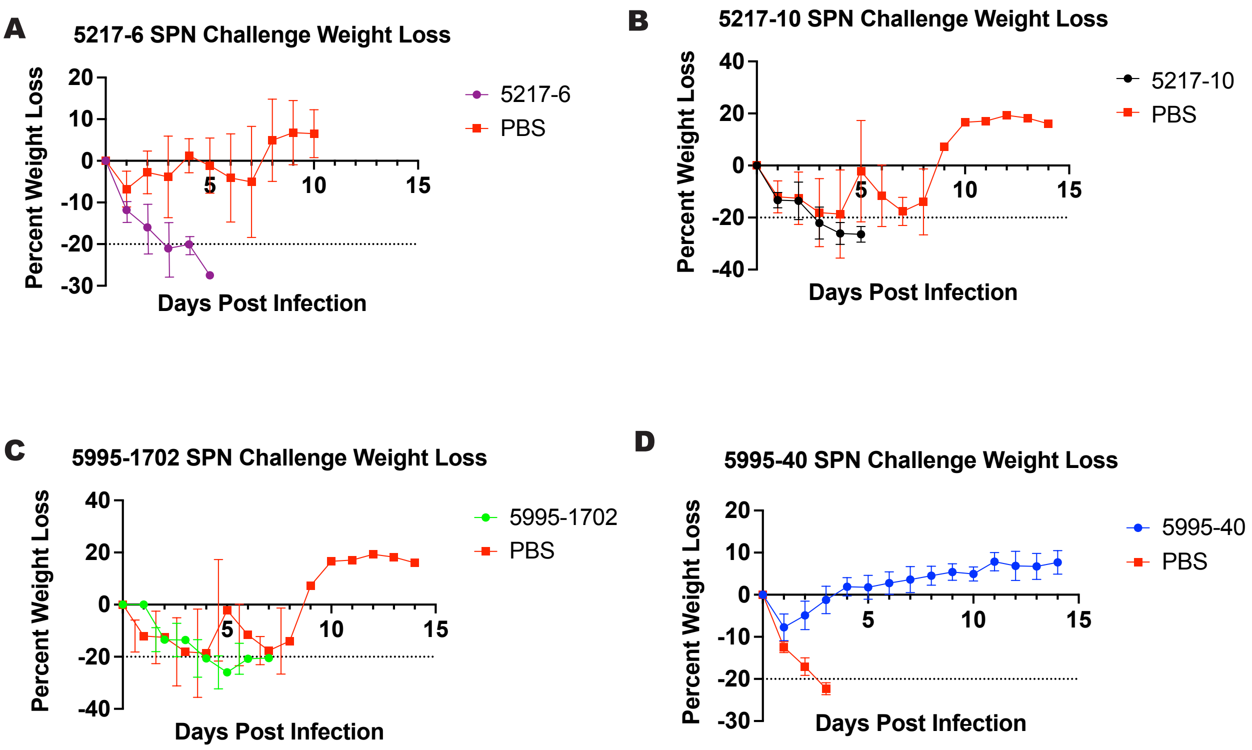


**Figure S4: Weight Loss Curves after WU2 infection.** Weight loss was measured over the course of 14 days after lethal WU2 infection, following a prophylactic treatment (15 mg/kg per mouse) of each respective mAb 2 hours prior to infection. Error bars are the standard deviation of weight loss of the living mice at each day post infection.

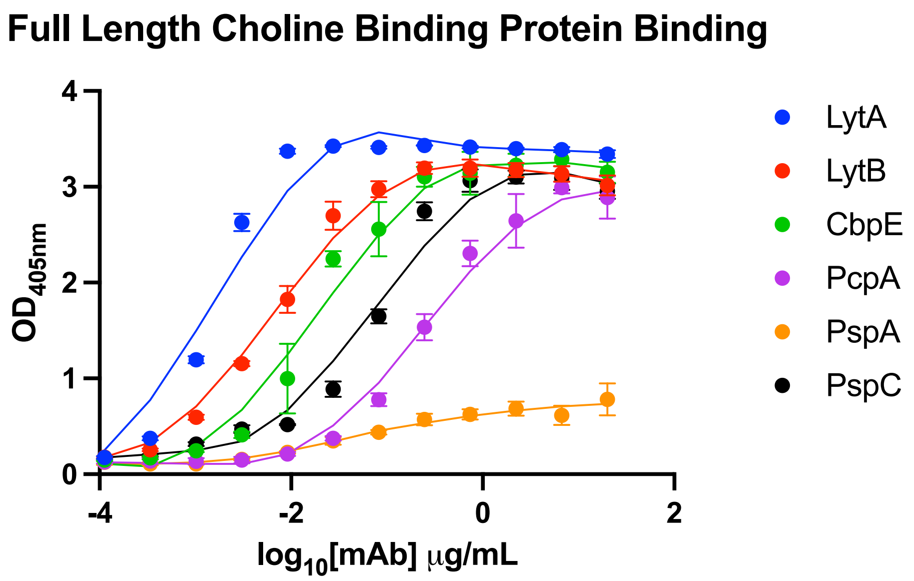


**Figure S5: ELISA binding of mAb 5995-40 to multiple full length choline binding proteins.** Binding of mAb 5995-40 to 6 choline-binding recombinant proteins by ELISA, with each point on the graph correlating to the $OD_{450}$ measured at that given mAb dilution. Error bars are the standard deviation of four technical replicates at each mAb dilution.

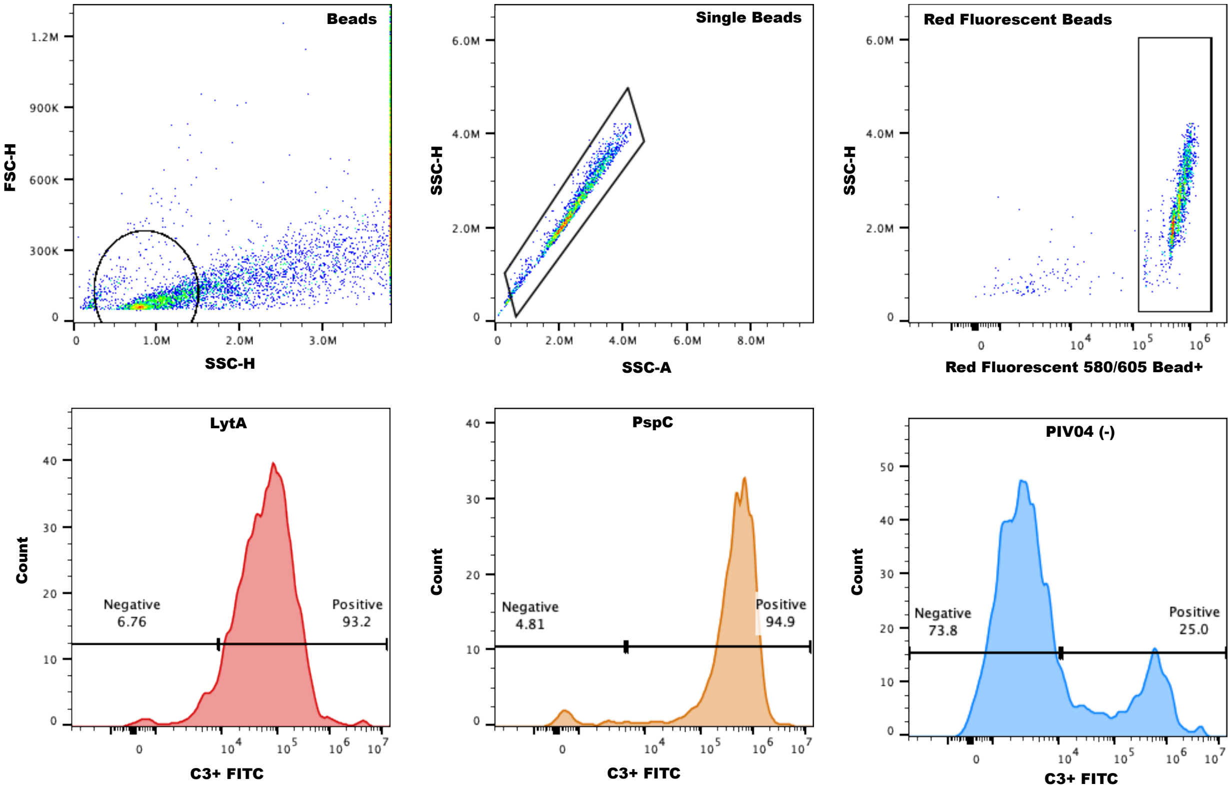


**Figure S6: Flow Panel to Determine C3 Complement Deposition Based on Antigen.** Red fluorescent beads were coated with antigen and incubated with mAb 5995-40 in the presence of guinea pig complement. Beads were then stained with an anti-C3 secondary antibody to detect C3 deposition. Using the gating strategy shown above, complement deposition was quantified as the percentage of $C3^{+}$ red fluorescent beads.

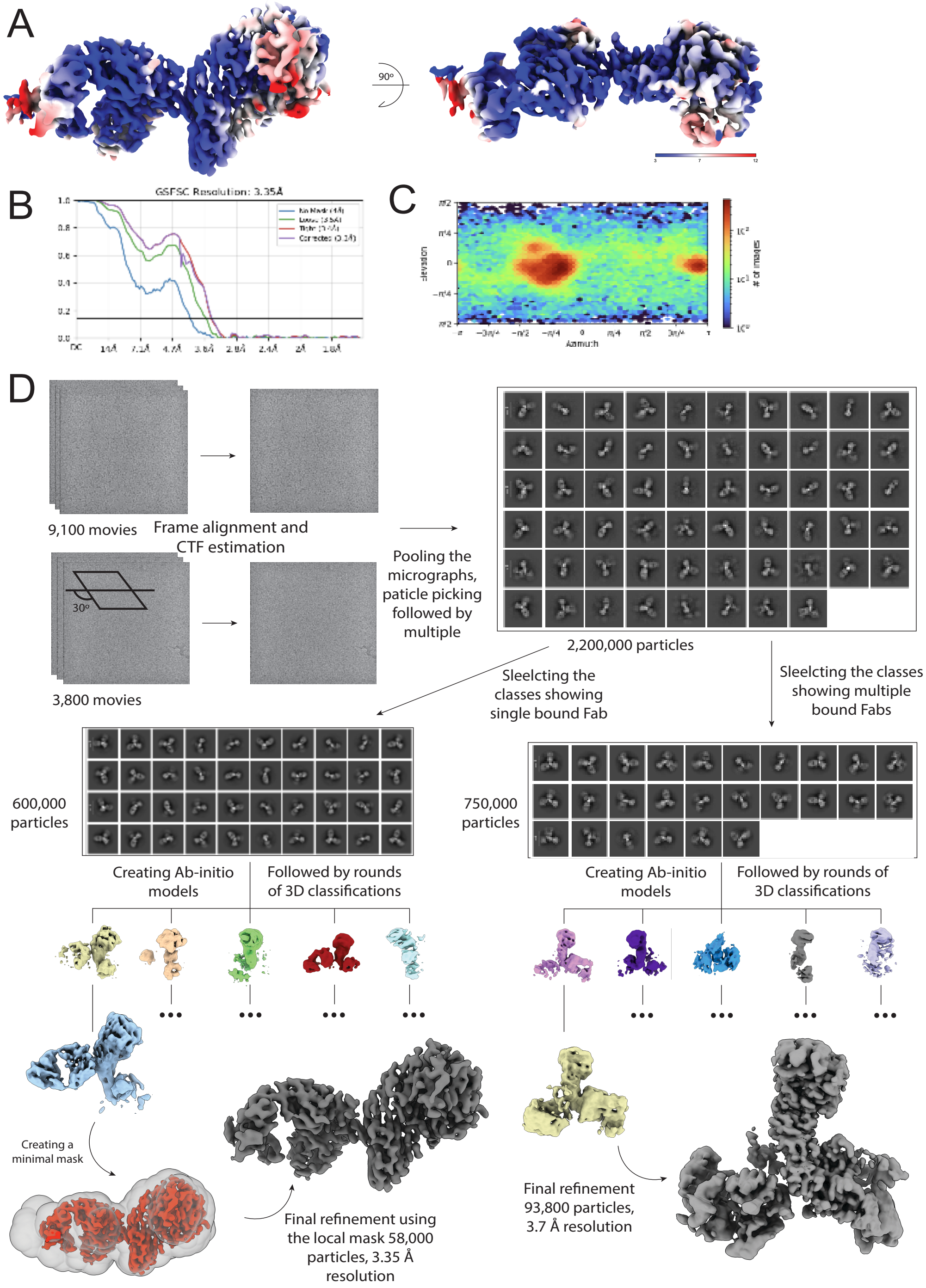


**Figure S7: CryoEM map quality and workflow.** A) Local resolution of the high-resolution map. B) GSFSC curve. C) Angular distribution of the particles. D) Overall workflow scheme, using two separate datasets (one tilted), 2D and 3D classifications and final result of two maps.

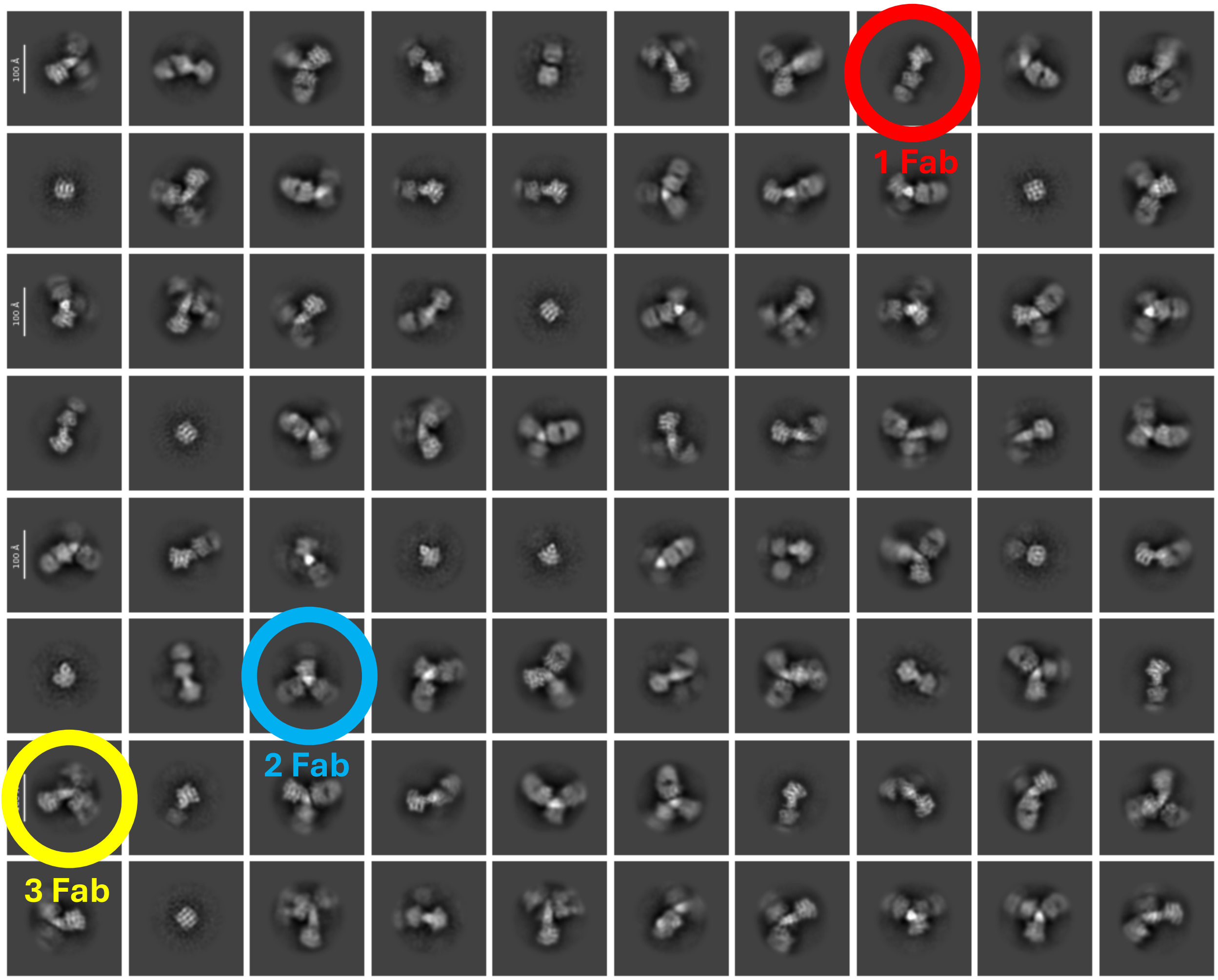


**Figure S8: CbpE complexed with Fab 5995-40 2D class averages.** 2D class averages from "small particle 2D classification" job in CryoSPARC showing the binding heterogeneity of 1-3 fabs in different distinguishable classes.
